# Lysine Demethylase 4A (KDM4A) Maintains Basal Body Architecture and Protects Against Ciliary Destabilization

**DOI:** 10.64898/2026.01.30.702865

**Authors:** Manga Motrapu, Lindsey Hudson, Pratim Chowdhury, Xiaoli Wang, Yuya Nakatani, Sofia Vargas-Hernandez, Sung Y. Jung, Menuka Karki, Cheryl Walker, Anna-Karin Gustavsson, Ruhee Dere

## Abstract

Primary cilia are sensory organelles essential for signaling and defects in formation, maintenance or structure underlie diverse ciliopathies. Here, we identify lysine demethylase 4A (KDM4A) as a previously unrecognized mediator of ciliogenesis. Using genetic depletion and pharmacologic inhibition, we show that KDM4A is required for cilia assembly and maintenance. Super-resolution imaging reveals KDM4A localization at the basal body, where it distinctively wraps around the centrioles. We uncover a direct interaction between KDM4A and Rootletin (CROCC), a structural protein mediating centriole cohesion, and demonstrate that *KDM4A* loss increases inter-centriolar distances, implicating basal body architecture in ciliary failure. Together, these findings define a demethylase-centrosome axis that integrates KDM4A activity with organelle biology, revealing new mechanisms underlying ciliogenesis.

## INTRODUCTION

Primary cilia are microtubule-based organelles that project from the cell surface during quiescence^1^. Cilia are found on virtually all mammalian cell types where they function to integrate diverse developmental and homeostatic pathways^2^. The ciliary membrane is enriched for sensory receptors and signaling molecules, enabling the transduction of signaling pathways including Hedgehog, WNT, and PDGFR which collectively regulate processes including differentiation, migration, and cell growth^3–5^. Consistent with this central role, mutations or loss of key ciliary proteins impair cilia function resulting in a broad spectrum of syndromic disorders collectively referred to as ciliopathies^6^.

Structurally, the ciliary axoneme (microtubule-based scaffold) emerges from the basal body, a modified centrosome that nucleates cilia formation^7^. Upon exit from mitosis, the mother centriole acquires specific distal and sub-distal appendages that permit initiation of axoneme extension^8^. Perturbations in centrosome numbers or architecture therefore frequently impair ciliogenesis^9^. Importantly, primary cilia are dynamic, built during quiescence when the cells no longer need to divide, and disassembled (resorbed) when the cells re-enter the cell cycle. Each of these processes, of ciliogenesis and cilia resorption, are governed by distinct sets of molecular effectors^7^.

Aurora kinase A (AURKA) is a well-characterized negative regulator of ciliogenesis. Activation of AURKA post-mitotically leads to the activation of histone deacetylase 6 (HDAC6) to promote ciliary disassembly through microtubule deacetylation, although AURKA could additionally have a direct impact on centrosome organization^10^. Elevated AURKA signaling in cells is associated with loss of primary cilia, whereas pharmacologic inhibition of its kinase activity results in a rescue of cilia in multiple cell types^11,12^.

Lysine demethylases are traditionally recognized for their roles in chromatin regulation^13^, DNA repair^14^ and genome stability^15^. Lysine demethylase 4A (KDM4A), a member of the Jumonji family of demethylases, removes di-and tri-methyl marks from histone H3 at lysine 36 (H3K36me2/3) and lysine 9 (H3K9me2/3)^16^. Emerging evidence, including work from our laboratory, indicates that KDM4A also localizes outside the nucleus^17,18^. We previously demonstrated that KDM4A associates with centrosomes during mitosis, where it localizes to the microtubule organizing centers (MTOCs) of the spindle^18^. Loss or inhibition of KDM4A induced numerical and structural centrosome abnormalities, driving genomic instability hinting at functions beyond chromatin regulation^18^. However, whether KDM4A contributes to basal body function and ciliogenesis remains unknown.

Here we demonstrate KDM4A is retained at the basal body during quiescence and is required for cilia formation and maintenance. Genetic depletion or pharmacologic inhibition of KDM4A abolishes ciliogenesis, an effect unique to KDM4A, as related demethylases fail to compensate. Enzymatic inhibition of KDM4A in cells that are pre-ciliated destabilizes existing cilia, underscoring a direct role for KDM4A at the basal body. In addition, pharmacologic inhibition of KDM4A reveals a threshold-dependent requirement for KDM4A enzymatic activity, supporting a chromatin-independent role at the basal body. KDM4A modulates AURKA levels, although AURKA inhibition does not rescue ciliogenesis in *KDM4A*-deficient cells indicating the involvement of additional mechanistic pathways. KDM4A inhibition increases inter-centriolar distances in cells lacking cilia, and immunoprecipitation-mass spectrometry (IP-MS) analysis identifies Rootletin (CROCC) as a KDM4A interactor, suggesting that KDM4A contributes to basal body architecture and spatial organization of centrioles. Together, these findings uncover an unexpected role for KDM4A in ciliogenesis and ciliary stability, expanding its functional repertoire beyond chromatin regulation, highlighting a previously unrecognized epigenetic-centrosome axis critical for cellular homeostasis.

## RESULTS

### KDM4A localizes to the basal body

We recently identified KDM4A as a centrosome-associated protein essential for preserving mitotic fidelity and genomic stability^18^. Beyond their canonical role as microtubule organizing centers (MTOCs) during mitosis, centrosomes serve an alternate, mutually exclusive function in quiescent cells as basal bodies that nucleate primary cilia^19^. Given that KDM4A localizes to centrosomes of the spindle during mitosis^18^, we investigated whether KDM4A could also localize to the basal body in non-dividing, ciliated cells.

In ciliated hTERT RPE-1 cells, diffraction-limited immunofluorescence microscopy revealed KDM4A enrichment at the base of the primary cilium, marked by acetylated α-tubulin, a ciliary axoneme marker (Figure 1A). Co-staining with γ-tubulin further confirmed KDM4A localization to the centrosome/basal body compartment (Figure 1B). Structured illumination microscopy (SIM) corroborated co-localization of KDM4A with centrosome-specific markers (γ-tubulin and Centrin-2) in ciliated cells (Figures S1A–S1D). To resolve sub-centrosomal distribution, we applied 3D single-molecule super-resolution imaging^20–25^, using direct stochastic optical reconstruction microscopy (dSTORM), which distinguished KDM4A at centrioles versus the pericentriolar matrix (Figures 1C-1E). 3D super-resolution reconstructions revealed that KDM4A preferentially associates and envelopes the proximal ends of the centrioles (Figures 1C-1F and S1E-S1G). Together, these data establish that KDM4A localizes to the basal body in ciliated cells, implicating this demethylase in basal body-associated functions.

**Figure 1.**
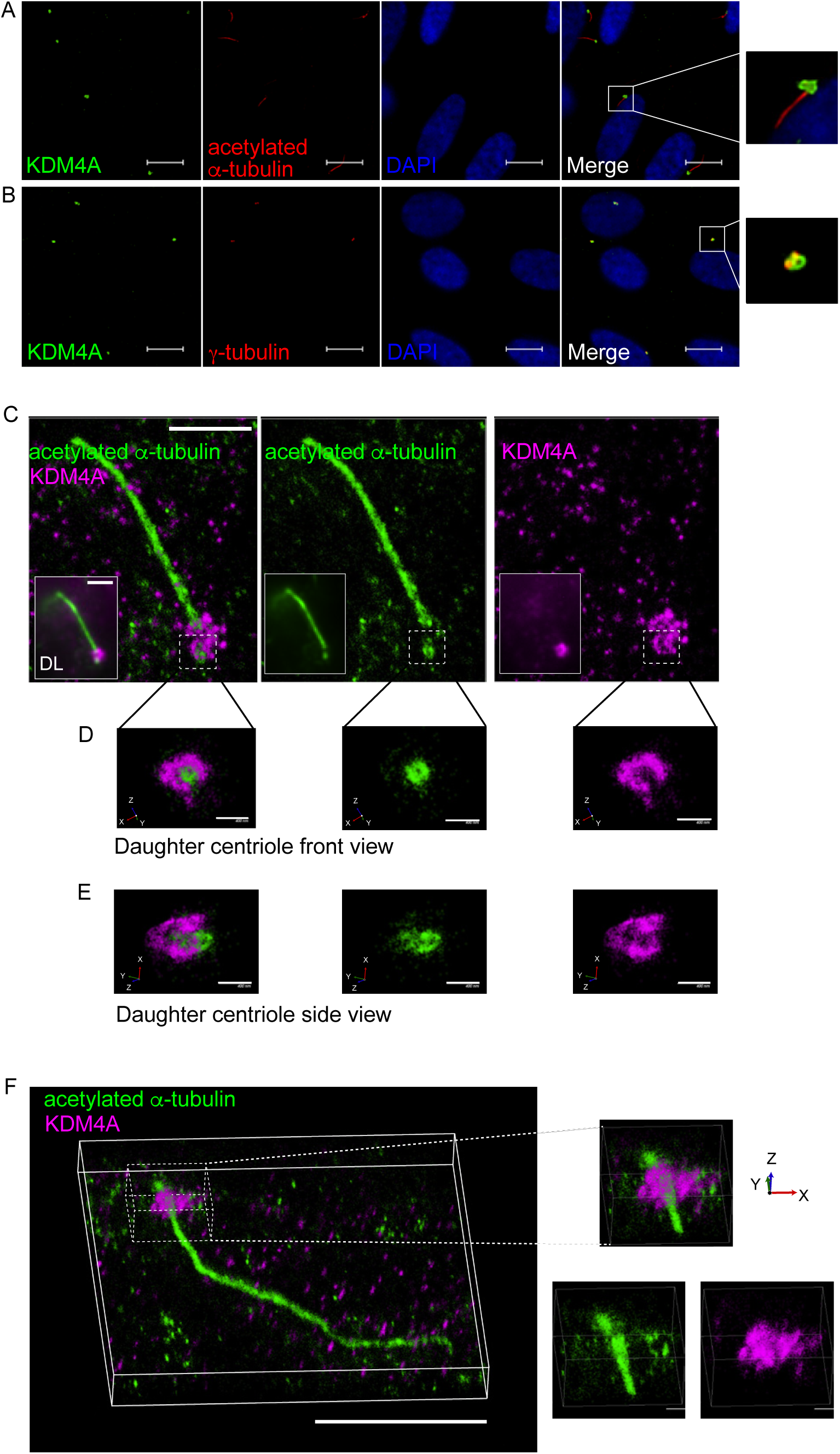
KDM4A localizes to the basal body. (A, B) Representative immunofluorescence images of hTERT RPE-1 (retinal pigmented epithelial) cells showing KDM4A (green) co-labeled with acetylated α-tubulin (red) (A) or γ-tubulin (B) (n=3). Nuclei were counterstained with DAPI (blue). Enlarged regions indicated by white boxes are shown to the right. Scale bar, 10 μm. (A) 3D super-resolution reconstruction of a cilium labeled for KDM4A (magenta) and acetylated α-tubulin (green), shown as merged and individual channels. Insets show corresponding diffraction-limited (DL) images. Scale bar, 3 µm. (D, E) Cross-sectional (D) and side (E) views of the daughter centriole within the same cilium. Scale bars, 400 nm. (F) 3D super-resolution reconstruction of another cilium labeled for KDM4A (magenta) and acetylated α-tubulin (green). Scale bar, 5 µm. Zoomed-in images of the proximal ciliary region are shown as merged (top, right) and individual channels (bottom, right). Scale bar, 1 µm. See also Figure S1.

### KDM4A is required for cilia formation

Having established KDM4A localization at the basal body, we next investigated its functional role in ciliogenesis. To assess whether KDM4A contributes to primary cilia formation, we acutely depleted *KDM4A* in hTERT RPE-1 cells using siRNA (siKDM4A) under serum starved conditions to induce ciliogenesis. Immunoblots confirmed efficient knockdown of *KDM4A* (Figure 2A). Immunofluorescence analysis 48 hours (h) post serum deprivation using acetylated α-tubulin (axoneme/cilia marker) and pericentrin (basal body marker) revealed a 2.6-fold reduction in the percentage of ciliated cells (Figures 2B and 2C). To exclude marker-specific effects i.e., cilia phenotype arising from altered acetylated α-tubulin levels, we used independent cilia markers – Arl13b (ADP-ribosylation factor-like protein 13b) and PGT (polyglutamylated tubulin). Consistent with the acetylated α-tubulin data, *KDM4A*-depletion resulted in a 2.7-fold (Figures 2D and 2E) and a 2.0-fold (Figures S2A-S2B) reduction with Arl13b and PGT, respectively. Next, we generated *KDM4A*-deficient hTERT RPE-1 cells using CRISPR-Cas9 mediated knockout (Figure 2F). Chronic loss of *KDM4A* (KDM4A CRISPR) significantly impaired ciliogenesis assessed using acetylated α-tubulin (Figures 2G and 2H) and Arl13b (Figure 2I), although the effect was less pronounced than acute depletion, suggesting adaptive mechanisms that could partially preserve cilia formation. Consistent with these data we did not observe any significant differences in cilia length using either ciliary marker (Figures S2C-S2D) in cells with chronic loss of *KDM4A*. Together, these results indicate that KDM4A is essential for efficient ciliogenesis, with an acute loss exerting a stronger disruptive effect than chronic deficiency.

**Figure 2.**
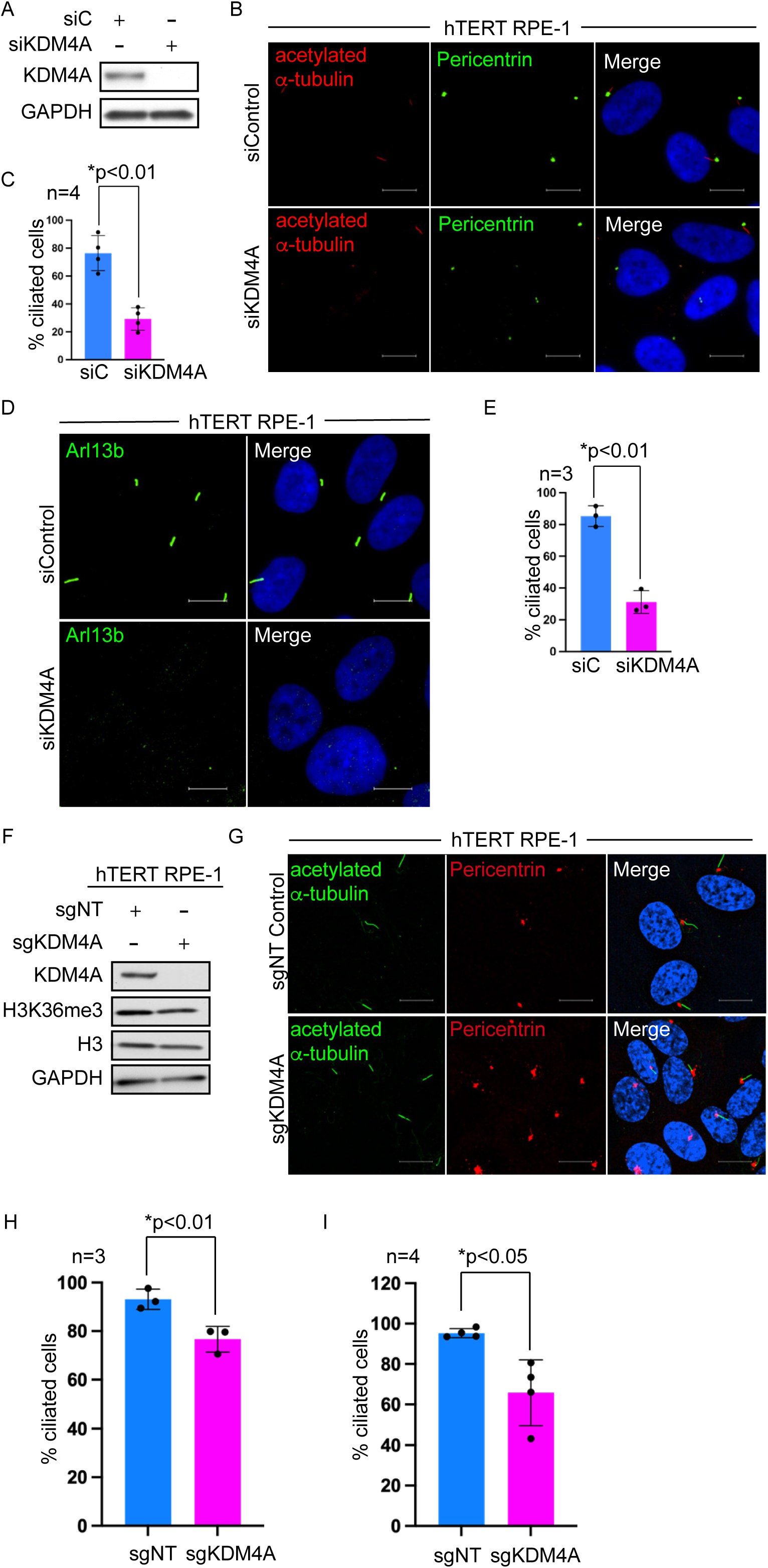
KDM4A is required for cilia formation. (A) Immunoblots of lysates from hTERT RPE-1 cells expressing siControl (non-targeting) or siKDM4A following serum starvation to induce ciliogenesis. (B, D) Representative immunofluorescence images of hTERT RPE-1 cells expressing siControl or siKDM4A (n=3) after 48 h of serum depletion, co-labeled with acetylated α-tubulin (cilia marker, red) and pericentrin (basal body marker, green) (B) or with Arl13b (cilia marker, green) (D). Nuclei were counterstained with DAPI (blue). Scale bar, 10 μm. (C, E) Quantification of ciliated hTERT RPE-1 cells (y-axis) expressing siControl (blue bars) or siKDM4A (pink bars) labeled with acetylated α-tubulin (C) or Arl13b (E). Each point represents values from individual experiments (n=4 for C; n=3 for E), with >500 cells analyzed per experiment. Data are shown as mean ± S.D.; * p-values as indicated (Welch’s t-test). (F) Immunoblots of lysates from CRISPR-generated *KDM4A* knockout (sgKDM4A) or non-targeting control (sgNT) hTERT RPE-1 cells. (G) Representative immunofluorescence images of sgNT and sgKDM4A hTERT RPE-1 cells co-labeled with acetylated α-tubulin (cilia marker, green) and pericentrin (basal body marker, red). Nuclei were counterstained with DAPI (blue). Scale bar, 10 μm. (H, I) Quantification of ciliated hTERT RPE-1 cells (y-axis) expressing sgNT (blue bars) and sgKDM4A (pink bars) labeled with acetylated α-tubulin (H) or Arl13b (I). Each point represents values from individual experiments (n=3 for H; n=4 for I), with >500 cells analyzed per experiment. Data are shown as mean ± S.D.; * p-values as indicated (Student’s t-test). See also Figure S2.

### Pharmacologic inhibition of KDM4A enzymatic activity impairs ciliogenesis

To complement genetic loss-of-function studies, we examined whether pharmacologic inhibition of KDM4A enzymatic activity affects ciliogenesis. KDM4A is a dioxygenase requiring iron and oxygen for its enzymatic function^26^. Treatment of hTERT RPE-1 cells with the iron chelator, deferoxamine (DFX, 250 μM), significantly reduced ciliogenesis resulting in a 1.7-fold decrease in ciliated cells upon quantification (Figures 3A and 3B).

**Figure 3.**
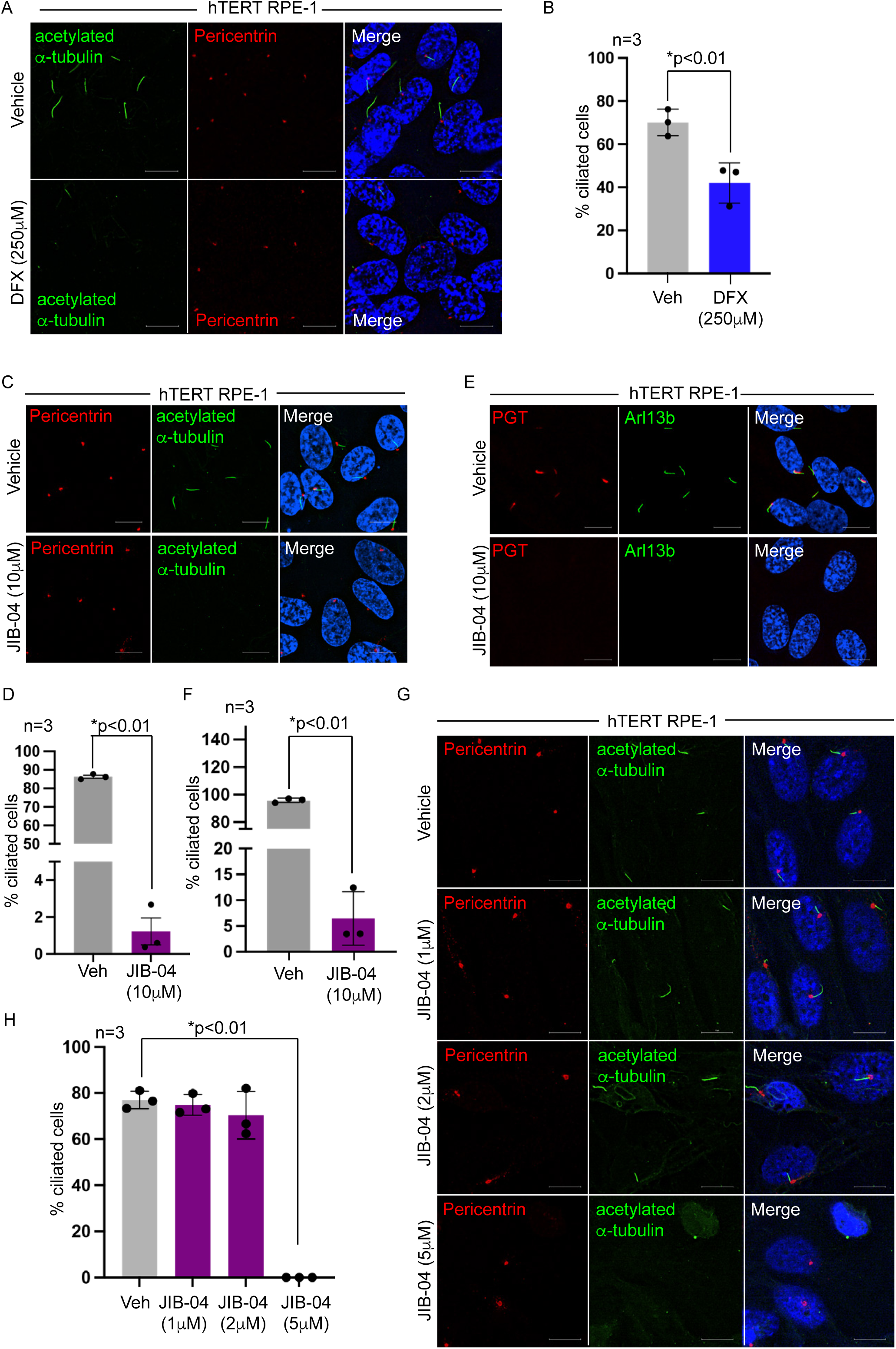
Pharmacologic inhibition of KDM4A enzymatic activity impairs ciliogenesis. (A) Representative immunofluorescence images of hTERT RPE-1 cells treated with vehicle (top) or Deferoxamine (DFX, bottom), co-labeled with acetylated α-tubulin (cilia marker, green) and pericentrin (basal body marker, red). Nuclei were counterstained with DAPI (blue). Scale bar, 10μm. (B) Quantification of ciliated hTERT RPE-1 cells (y-axis) treated with vehicle (gray bars) and DFX (blue bars). Each point represents values from individual experiments (n=3), with >500 cells analyzed per experiment. Data are shown as mean ± S.D.; * p-values as indicated (Student’s t-test). (C, E) Representative immunofluorescence images of hTERT RPE-1 cells treated with vehicle (top) or JIB-04 (10 μm, bottom), co-labeled with acetylated α-tubulin (cilia marker, green) and pericentrin (basal body marker, red) (C) or Arl13b (cilia marker, green) and PGT (cilia marker, red) (E). Nuclei were counterstained with DAPI (blue). Scale bar, 10 μm. (D, F) Quantification of ciliated hTERT RPE-1 cells (y-axis) treated with vehicle (gray bars) or JIB-04 (plum bars) labeled with acetylated α-tubulin (D) or Arl13b (F). Each point represents values from individual experiments (n=3 for each), with >500 cells analyzed per experiment. Data are shown as mean ± S.D.; * p-values as indicated (Welch’s t-test). (G) Representative immunofluorescence images of hTERT RPE-1 cells treated with vehicle (top) or increasing doses of JIB-04 (bottom), co-labeled with acetylated α-tubulin (cilia marker, green) and pericentrin (basal body marker, red). Nuclei were counterstained with DAPI (blue). Scale bar, 10 μm. (H) Quantification of ciliated hTERT RPE-1 cells (y-axis) treated with vehicle (gray bars) or JIB-04 (plum bars). Each point represents values from individual experiments (n=3), with >500 cells analyzed per experiment. Data are shown as mean ± S.D.; * p-values as indicated (Welch’s t-test). See also Figure S2.

To specifically target KDM4A’s demethylase activity, we used a pan-Jumonji demethylase inhibitor, JIB-04^27^. Cells treated with 10 μM JIB-04 at the onset of serum deprivation (to induce cilia formation) resulted in a dramatic 45-fold reduction in ciliogenesis as assessed by acetylated α-tubulin staining (Figures 3C-3D). The few cilia that formed were markedly shorter (Figure S2E), with Arl13b staining further corroborating both the reduction in cilia frequency (Figures 3E and 3F) and the shortening of cilia length in JIB-04 treated cells (Figure S2F). Dose-response analysis revealed minimal effects at lower concentrations with pronounced loss of cilia emerging at the 5 μM JIB-04 dose (Figures 3G and 3H). These findings indicate that KDM4A enzymatic function is a critical determinant of ciliogenesis.

### KDM4A exclusively impairs ciliogenesis among Jumonji demethylases targeted by JIB-04

Given that chromatin modifying enzymes often exhibit redundancy in their histone-related functions^28^, we investigated whether other Jumonji family demethylases targeted by JIB-04 influence cilia formation. In addition to KDM4A, JIB-04 targets four other Jumonji demethylases including KDM4B, KDM4C, KDM5A and KDM6B^27^. To assess their individual contributions to ciliogenesis, we systematically knocked down each of these demethylases in hTERT RPE-1 cells (Figures 4A-4D) and evaluated cilia formation using acetylated α-tubulin staining. Knockdown of *KDM4B*, *KDM4C*, *KDM5A* and *KDM6B* (via siRNA) did not significantly affect cilia formation, which remained comparable to the siControl-treated cells (Figures 4E and 4F). These findings were corroborated by Arl13b (Figures 4G and 4H) and PGT staining (Figures S3A and S3B), confirming that cilia formation was unaffected in cells depleted of these enzymes. Thus, these results indicate that KDM4A plays a unique and non-redundant role in regulating ciliogenesis.

**Figure 4.**
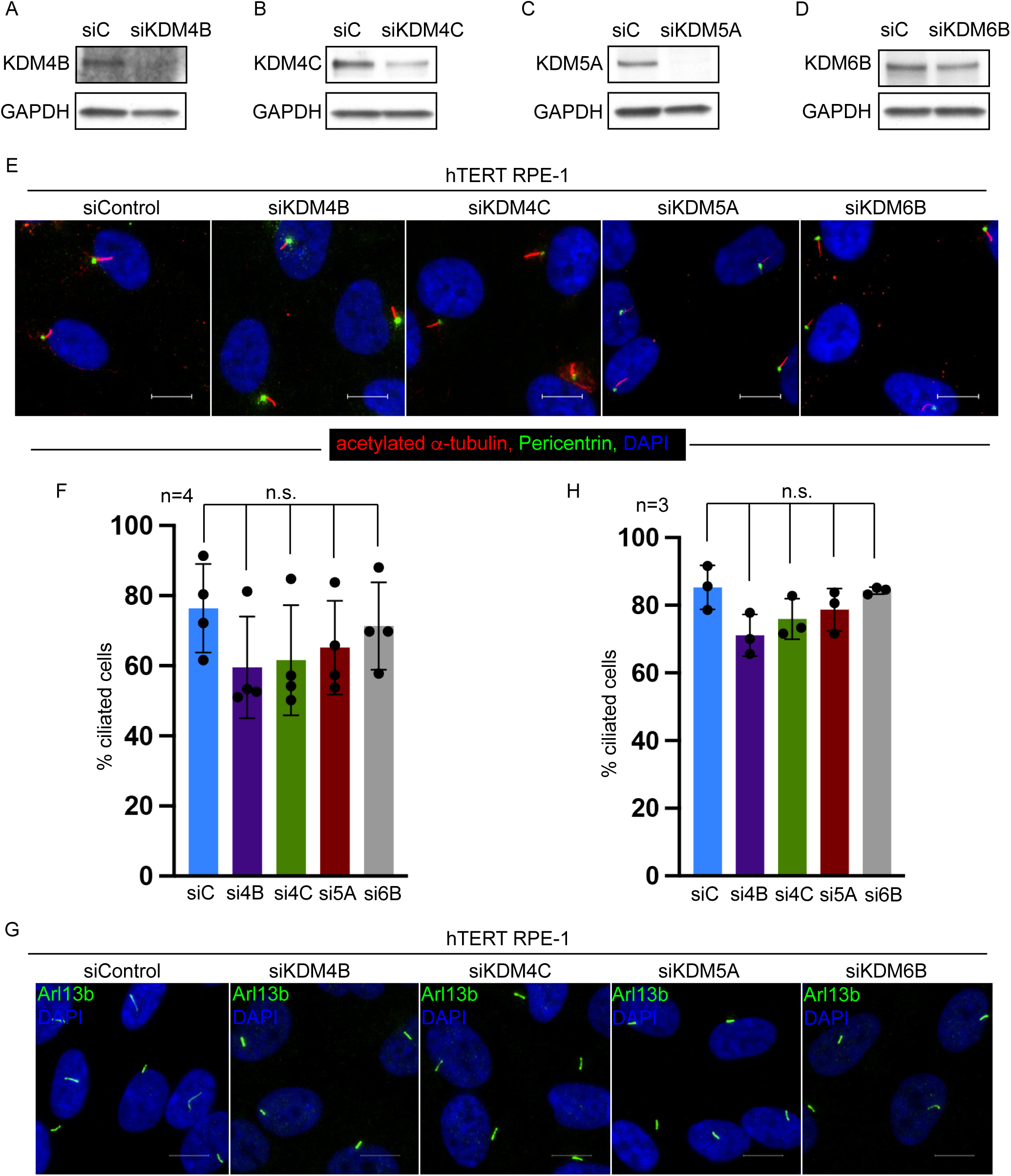
KDM4A exclusively impairs ciliogenesis among Jumonji demethylases targeted by JIB-04. (A-D) Immunoblots of lysates from hTERT RPE1 cells expressing siControl (non-targeting) or siRNA’s targeting *KDM4B* (A), *KDM4C* (B), *KDM5A* (C) and *KDM6B* (D) following serum starvation to induce ciliogenesis. (E, G) Representative immunofluorescence images of hTERT RPE-1 cells expressing siControl or siKDM4B, siKDM4C, siKDM5A and siKDM6B (n=3) after 48 h of serum depletion, co-labeled with acetylated α-tubulin (cilia marker, red) and pericentrin (basal body marker, green) (E) or Arl13b (cilia marker, green) (G). Nuclei were counterstained with DAPI (blue). Scale bar, 10 μm. (F, H) Quantification of ciliated hTERT RPE-1 cells (y-axis) expressing siControl (blue bars) or siKDM4B (purple bars), KDM4C (green bars), KDM5A (maroon bars) and KDM6B (gray bars), labeled with acetylated α-tubulin (F) or Arl13b (H). Each point represents values from individual experiments (n=4 for F; n=3 for H), with >500 cells analyzed per experiment. Data are shown as mean ± S.D.; * p-values as indicated (Student’s t-test). See also Figure S3.

### KDM4A enzymatic activity preserves cilia stability

Having established a functional role for KDM4A in cilia formation, we next examined its contribution to cilia maintenance. To assess cilia stability, hTERT RPE-1 cells were first serum-starved to induce cilia formation, prior to treatment with JIB-04 to inhibit KDM4A demethylase activity. A 24 h treatment with JIB-04 resulted in a marked reduction in the percentage of ciliated cells compared to cells treated with an inactive Z-isomer of JIB-04 (Figures 5A and 5B). Dose-response analysis revealed that 5 μM and 10 μM JIB-04 (Figures S4A and S4B) were sufficient to induce cilia loss in pre-ciliated cells, consistent with concentrations that impaired cilia formation (Figures 3G and 3H).

**Figure 5.**
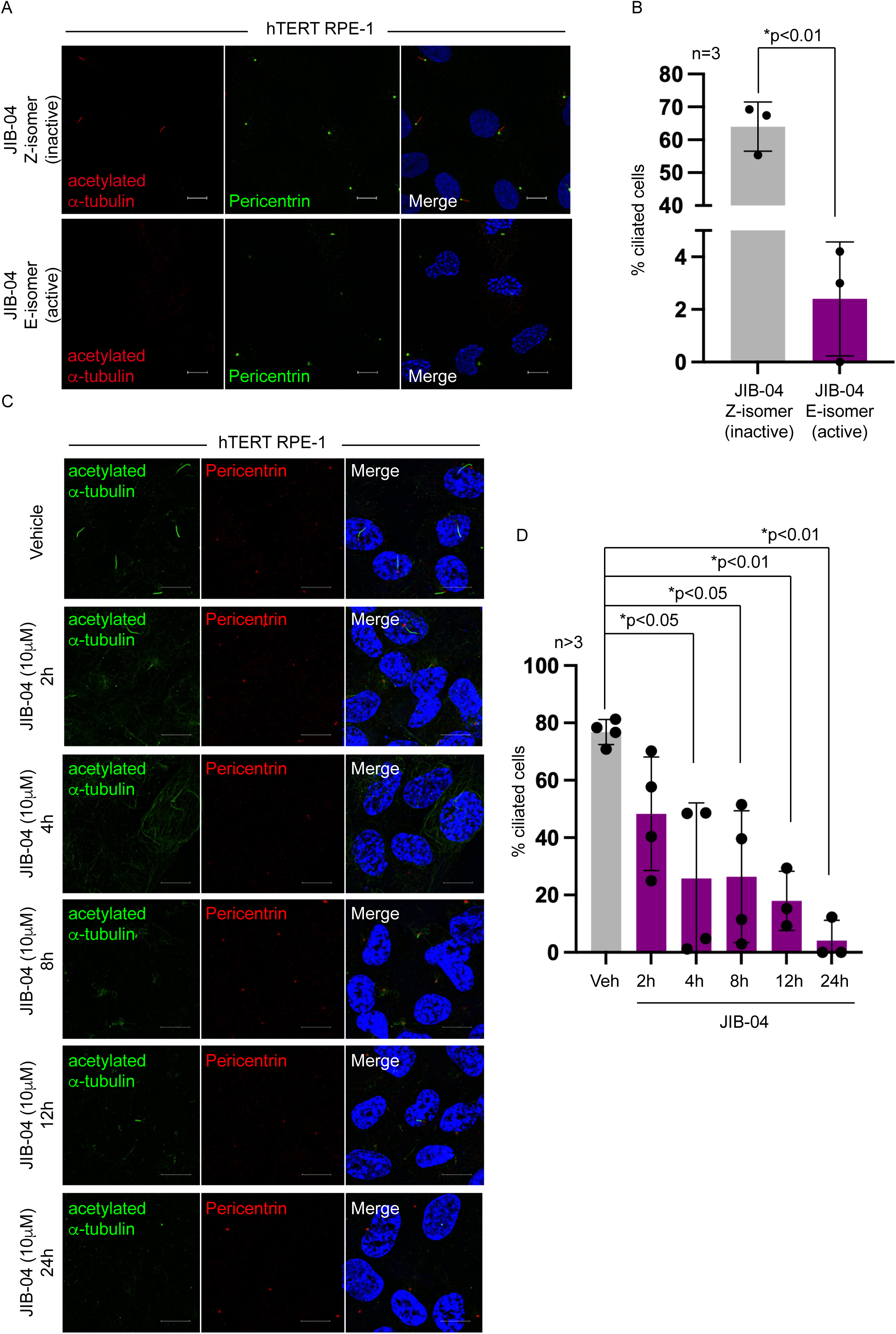
KDM4A enzymatic activity preserves cilia stability. (A) Representative immunofluorescence images of pre-ciliated hTERT RPE-1 cells treated for 24 h with JIB-04 Z-isomer (inactive compound as control, top) or JIB-04 E-isomer (active compound, bottom), co-labeled with acetylated α-tubulin (cilia marker, red) and pericentrin (basal body marker, green). Nuclei were counterstained with DAPI (blue). Scale bar, 10 μm. (B) Quantification of pre-ciliated hTERT RPE-1 cells (y-axis) treated with JIB-04 Z-isomer (gray bar) or E-isomer (plum bar) for 24h. Each point represents values from individual experiments (n=3), with >500 cells analyzed per experiment. Data are shown as mean ± S.D.; * p-values as indicated (Student’s t-test). (C) Representative immunofluorescence images of pre-ciliated hTERT RPE-1 cells treated with vehicle or JIB-04 (10μM) for the indicated time points (2h, 4h, 8h, 12h, 24h), co-labeled with acetylated α-tubulin (cilia marker, green) and pericentrin (basal body marker, red). Nuclei were counterstained with DAPI (blue). Scale bar, 10 μm. (D) Quantification of pre-ciliated hTERT RPE-1 cells (y-axis) treated with vehicle (gray bar) or JIB-04 (plum bars) for 2h, 4h, 8h, 12h and 24h. Each point represents values from individual experiments (n=4 for vehicle, 2h, 4h and 8h; n=3 for 12h and 24h), with >500 cells analyzed per experiment. Data are shown as mean ± S.D.; * p-values as indicated (Welch’s t-test). See also Figures S4 and S5.

To determine the kinetics of cilia disassembly following KDM4A inhibition, we performed a time-course analysis with JIB-04 treatment for 2 h, 4 h, 8 h, 12 h and 24 h (Figures 5C and 5D). A significant decrease in cilia was observed as early as 4 h post-treatment with progressive loss at the later time points (Figure 5D). Live cell imaging of hTERT RPE-1 cells expressing GFP-Arl13b confirmed rapid cilia shortening upon JIB-04 exposure (Figures S5A and S5B), further validating the destabilizing effect of KDM4A on primary cilia. Collectively, these data demonstrate that KDM4A contributes to both cilia formation and stability, establishing its enzymatic activity as a critical determinant of cilia integrity.

### AURKA levels are modulated by KDM4A inhibition

Given that elevated Aurora kinase A (AURKA) expression is strongly associated with impaired ciliogenesis^10^, we evaluated whether KDM4A inhibition influenced AURKA levels. We treated hTERT RPE-1 cells with JIB-04 at the time of serum depletion to induce ciliation and AURKA expression was assessed after 48 h. JIB-04 treatment led to a marked increase in AURKA protein levels (Figures 6A-6B), correlating with the observed failure to form cilia (Figures 3C-3F). We next asked whether AURKA upregulation also occurred during cilia destabilization associated with KDM4A inhibition (Figure 5). In pre-ciliated cells treated with JIB-04, AURKA expression similarly increased (Figures 6C-6D) at early time points, suggesting that KDM4A may influence AURKA levels during both cilia formation and maintenance. To determine if this increase reflected transcriptional regulation, we measured AURKA transcript levels, which showed no significant differences in mRNA abundance (Figure 6E), indicating that KDM4A inhibition elevates AURKA protein through a post-transcriptional mechanism. However, co-inhibition of both KDM4A (JIB-04, 5 μM) and AURKA (alisertib, 2 μM) failed to rescue ciliation (Figure S6A). Similarly, co-inhibition of AURKA (alisertib, 2 μM) during cilia destabilization using JIB-04 (10 μM) also failed to preserve cilia (Figure S6B), indicating that AURKA upregulation alone does not fully account for the ciliary phenotype.

**Figure 6.**
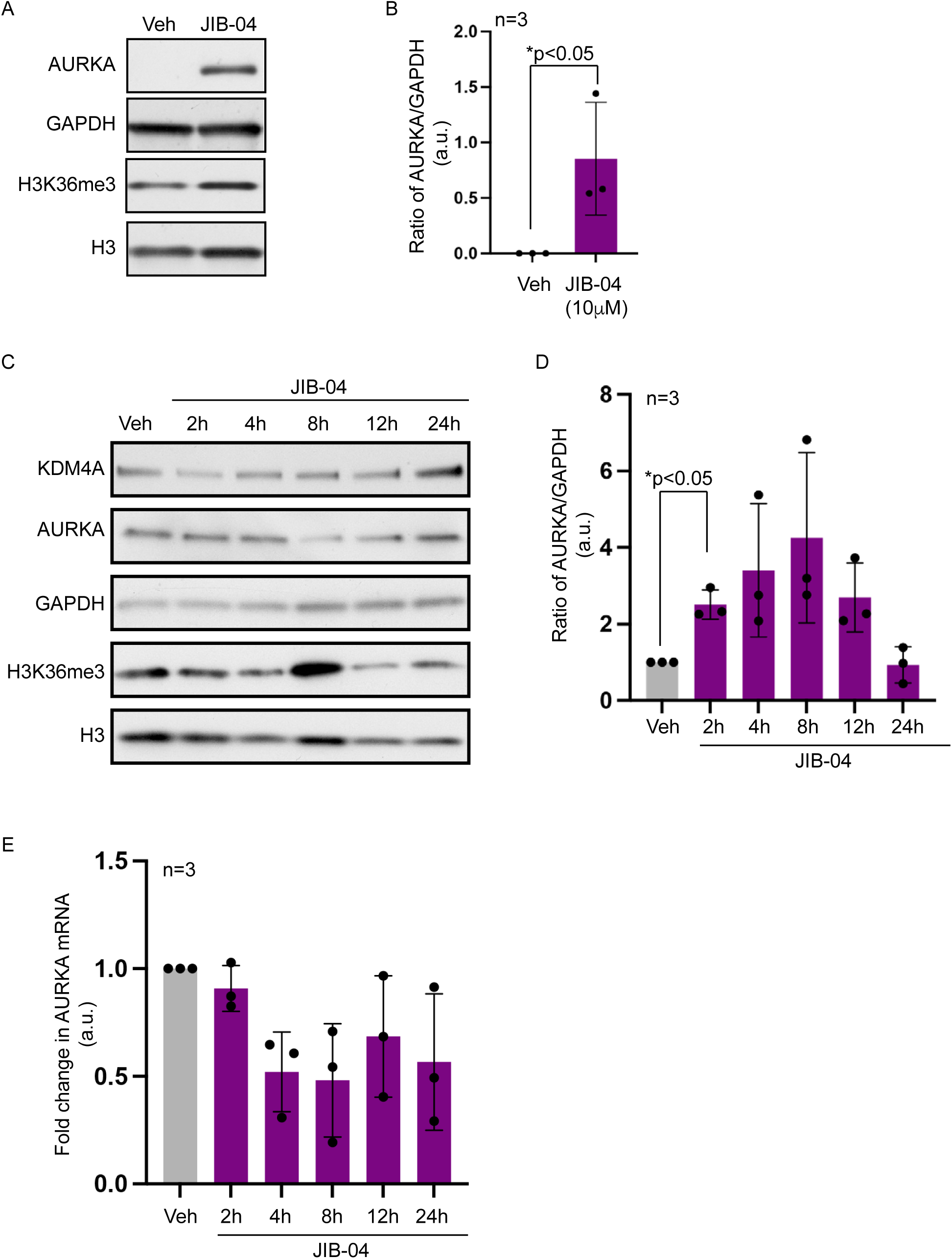
AURKA levels are modulated by KDM4A inhibition. (A) Immunoblots of lysates from hTERT RPE-1 cells treated with vehicle or JIB-04 during cilia formation. (B) Quantification of densitometric analysis of AURKA immunoblots normalized to GAPDH from vehicle treated (gray bar) and JIB-04 treated (plum bar) hTERT RPE-1 cells. Each point represents values from individual experiments (n=3). Data are shown as mean ± S.D.; * p-values as indicated (Student’s t-test). (C) Immunoblots of lysates from pre-ciliated hTERT RPE-1 cells treated with vehicle or JIB-04 for the indicated times (2 h, 4 h, 8 h, 12 h, and 24 h). (D) Quantification of densitometric analysis of AURKA immunoblots normalized to GAPDH from vehicle-treated (gray bar) and JIB-04-treated (plum bars) HTERT RPE-1 cells at 2 h, 4 h, 8 h, 12 h and 24 h. Each point represents values from individual experiments (n=3). Data are shown as mean ± S.D.; * p-values as indicated (Student’s t-test). (E) RT-PCR analysis showing fold change in AURKA mRNA in JIB-04 treated (2 h, 4 h, 8 h, 12 h, and 24 h; plum bars) relative to vehicle control (gray bar). Each point represents values from individual experiments (n=3). Data are shown as mean ± S.D.; * p-values as indicated (Student’s t-test). See also Figure S6.

### KDM4A interacts with Rootletin (CROCC) to regulate inter-centriolar distances

JIB-04 treated cells exhibited increased inter-centriolar distances between centrin-2 puncta compared to vehicle controls (Figures 7A-7B and Figures S7A-S7B)), suggesting impaired centriole cohesion. Notably, this effect was variable with nearly 80% of JIB-04 treated cells displaying above average inter-centriolar distances compared to the vehicle treated cells. To identify potential KDM4A interactors independent of AURKA regulation, we performed immunoprecipitation-mass spectrometry (IP-MS). Several centrosome-associated proteins were significantly enriched (Figure 7C), including Pericentrin (PCNT) and Rootletin (CROCC). Given that KDM4A localizes to the proximal ends of centrioles (Figure 1C) where Rootletin resides^29^, we tested for a direct interaction. Co-immunoprecipitation confirmed KDM4A binding with Rootletin (Figure 7D) in hTERT RPE-1 cells independent of JIB-04 treatment. Immunofluorescence imaging further validated the co-localization of KDM4A and Rootletin at the basal body (Figure 7E). These findings support a model in which KDM4A likely maintains basal body architecture through Rootletin-mediated centriole cohesion, and its inhibition disrupts this interaction, thereby destabilizing the ciliary apparatus.

**Figure 7.**
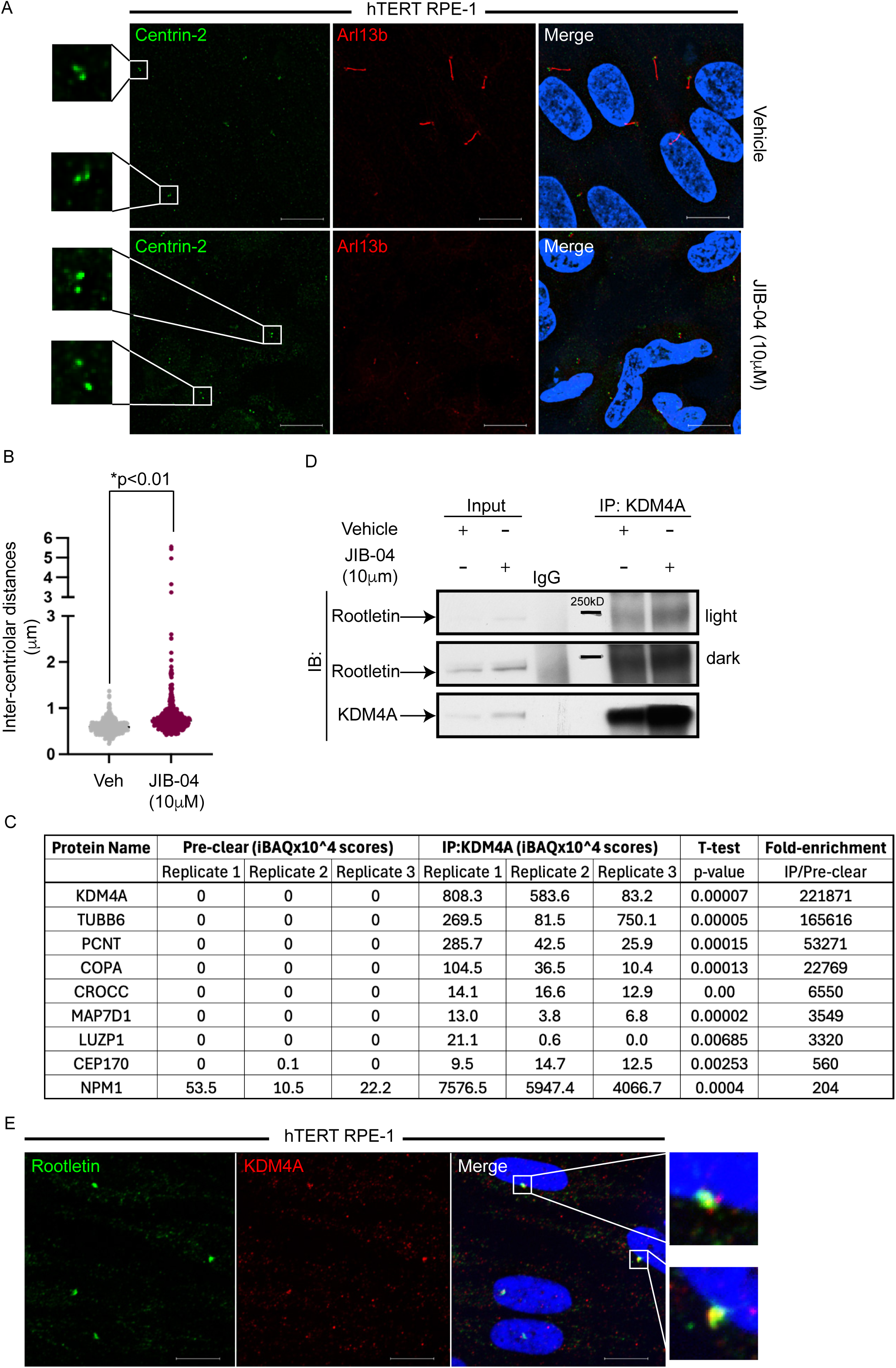
KDM4A interacts with Rootletin (CROCC) to regulate inter-centriolar distances. (A) Representative immunofluorescence images of pre-ciliated hTERT RPE-1 cells treated with vehicle (top) or JIB-04 (bottom), co-labeled with Centrin-2 (centriole marker, green) and Arl13b (cilia marker, red), n=3. Nuclei were counterstained with DAPI (blue). Enlarged regions indicated by white boxes are shown to the left. Scale bar, 10 μm. (B) Quantification of inter-centriolar distances (μm, y-axis) in pre-ciliated hTERT RPE-1 cells treated with vehicle (gray) or JIB-04 (plum) for 24h. Each point represents values from one representative experiment, with >400 cells analyzed. * p-values as indicated (Welch’s t-test). (C) Table summarizing immunoprecipitation mass spectrometry (IP-MS) analysis showing iBAQ scores from three replicates of immunoprecipitation with anti-KDM4A antibody. The fold enrichment of IP/pre-clear is shown in the last column, with p-values indicated in the penultimate column. (D) Co-immunoprecipitation analysis demonstrating KDM4A pulldown of Rootletin from hTERT RPE-1 cells treated with vehicle or JIB-04. (E) Representative immunofluorescence images of hTERT RPE-1 cells co-labeled with KDM4A (red) and Rootletin (green), n=3. Nuclei were counterstained with DAPI (blue). Enlarged regions indicated by white boxes are shown to the right. Scale bar, 10 μm. See also Figure S7.

## DISCUSSION

Our study identifies KDM4A as a previously unrecognized regulator of primary cilia, demonstrating its localization at the basal body and its essential role in both cilia formation and the maintenance of cilia stability. Using complementary genetic and pharmacologic approaches, we show that KDM4A is required for ciliogenesis and that its loss in ciliated cells destabilizes existing cilia. These findings extend our recent work establishing KDM4A as a centrosome-associated protein during mitosis, where its enzymatic activity preserves the integrity of the spindle pole and centrosome architecture^18^. Here we expand this centrosome-specific role of KDM4A to the basal body, a modified centrosome, establishing the importance of this epigenetic demethylase in regulating extranuclear organelle biology.

A striking observation in our study is the phenotypic difference between acute and chronic loss of *KDM4A*. Acute depletion of *KDM4A* either genetically or via pharmacologic inhibition resulted in robust loss of cilia, whereas chronic loss of *KDM4A* produced a milder phenotype. This discrepancy suggests activation of adaptative, or compensatory mechanisms triggered in cells with long-term *KDM4A*-deficiency. Such compensation could involve upregulation of other redundant demethylases, or transcriptional reprogramming of genes, consistent with the known functional redundancy among chromatin modifiers^30^. However, knockdown of the other members of the Jumanji family of demethylases, including *KDM4B*, *KDM4C*, *KDM5A* and *KDM6B*, targeted by the pan-Jumonji inhibitor^31^, JIB-04, did not significant impact ciliogenesis underscoring the specificity of KDM4A in regulating cilia. A similar temporal adaptation to chronic loss of *KDM4A* was observed in our prior studies where the severity of mitotic defects associated with *KDM4A* depletion diminished over time in culture^18^.

Pharmacologic studies further support a threshold-dependent requirement for KDM4A enzymatic activity in ciliogenesis. Inhibition of Jumonji demethylases with JIB-04 disrupted ciliogenesis only at concentrations greater than 5 μM, indicating that ciliogenesis is buffered against partial loss of demethylase activity and collapses when a critical enzymatic threshold is exceeded. Although nuclear functions of KDM4-family demethylases are inhibited at sub-micromolar concentrations^31^, the suppression of ciliogenesis at higher JIB-04 concentrations likely reflects differences in biochemical potency and effective target engagement within the cellular context of the basal body. Factors such as enzyme abundance, compartmental accessibility, and protein-protein interactions may shift these functional requirements. Importantly, this observation provides evidence for a chromatin-independent role of KDM4A in ciliogenesis, as nuclear KDM4A activity would have been efficiently suppressed at lower concentrations.

AURKA a known driver of cilia disassembly^10^ and oncogenesis^32^ is elevated in models of impaired ciliogenesis^33^. We found that KDM4A inhibition during ciliogenesis and in pre-ciliated cells resulted in elevated AURKA protein abundance, suggesting that KDM4A modulates AURKA levels. However, co-inhibition of AURKA (using a pharmacologic inhibitor, alisertib^34^) in *KDM4A*-depleted cells did not rescue ciliogenesis, indicating that AURKA upregulation alone does not fully account for the observed phenotype.

Given that AURKA elevation is especially pronounced during cilia assembly but not during cilia maintenance, our data support a model in which KDM4A functions as a gatekeeper of cilium assembly, whereas ciliary stability, may depend on a distinct Rootletin-dependent axis. KDM4A regulation of AURKA appears independent of transcriptional control, suggesting post-translational mechanisms, although whether KDM4A directly demethylates AURKA or influences stability or levels through intermediary factors remains to be determined. This link between KDM4A and AURKA provides a compelling framework for understanding how epigenetic dysregulation can drive cilia loss and contribute to oncogenic signaling.

Our findings reveal a second mechanism beyond AURKA regulation in which KDM4A supports ciliogenesis by interacting with the structural protein Rootletin to maintain centriole cohesion. Super-resolution imaging shows that KDM4A localizes at the basal body in a distinctive ‘French hot-dog’ like wrapping pattern around the centrioles, particularly at the mother-daughter interface suggesting a structural role in basal body organization. Consistent with this, we demonstrate that KDM4A physically associates with Rootletin (CROCC), a core component of the fibrous linkages that stabilize the centriole pairs^35,36^ and that KDM4A depletion increases inter-centriolar distances, indicating compromised cohesion. Although Rootletin is dispensable for normal cilia assembly in Drosophila^29^, its loss disrupts centriole cohesion^37^ and impairs cilia stability^38^ in mammals and *C. elegans*^39^, supporting a model in which Rootletin maintains ciliary integrity rather than initiating ciliogenesis. Thus, defective Rootletin-dependent structural support likely contributes to the ciliary defects in *KDM4A*-deficient cells. Given the emerging interest in phase separation^40^ and fiber-based cohesion at the centrosome^41,42^, future studies should determine whether KDM4A regulates Rootletin assembly or stability through demethylation-dependent mechanisms. This possibility aligns with our previous observations of centrosome fragmentation and cohesion defects in *KDM4A*-deficient mitotic cells^18^.

Our identification of KDM4A as a basal body associated demethylase whose enzymatic function regulates ciliogenesis, suggests the existence of other epigenetic modifiers in operation at the cilium. Recent methylome studies in *Chlamydomonas* identified over a hundred methylated ciliary proteins^43^ supporting the idea that methylation/demethylation is integral to ciliary function. Together, our findings define an unrecognized demethylase-centrosome axis that integrates chromatin modifiers with organelle architecture and signaling. Future studies should define the methylome of ciliary proteins and elucidate how *KDM4A*-dependent demethylation shapes basal body structure, protein recruitment and signaling, while identifying methyltransferases that establish these methyl marks. These insights may uncover novel therapeutic opportunities for ciliopathies and cancers characterized by ciliary dysfunction.

## RESOURCE AVAILABILITY

Requests for further information and resources should be directed to and will be fulfilled by the lead contact, Ruhee Dere.

### Material availability

All unique/stable reagents generated in this study are available from the lead contact (see above) with a completed institutional materials transfer agreement (MTA).

### Data availability

All data reported in this paper will be shared by the lead contact upon request.

## Supporting information

Manuscript

## ACKNOWLEDGEMENTS

The authors would like to thank the members of the Dere lab for their thoughtful comments and feedback. This work was supported by research funding from the Cancer Prevention Research Institute of Texas (RP220332), Department of Defense CDMRP KCRP (KC210123), NIH/NCI (R01-CA275082) to RD; Cancer Prevention and Research Institute of Texas (RR200035), and the NIH/NIGMS (R35GM155365) to AKG; a SynthX Award to RD and AKG; the Rice University Provost’s TMC Collaborator Seed Fund Program (PTC2401) to AKG and RD; NIH/NCI (R35CA231993) and NIH/NIEHS (P30ES030285) to CLW.

## AUTHOR CONTRIBUTIONS

MM Data curation, investigation, methodology, project administration, validation, visualization, writing

LH Investigation, methodology, validation, visualization, writing PC Investigation, methodology, validation

XW Investigation, methodology, validation

YN Investigation, methodology, visualization, writing SVH Investigation, methodology

SYJ Investigation, methodology, validation MK Investigation, methodology, validation

CLW Conceptualization, resources, funding acquisition, editing

AKG Conceptualization, formal analysis, resources, funding acquisition, supervision, editing

RD Conceptualization, data curation, investigation, formal analysis, project administration, funding acquisition, supervision, writing and editing

## DECLARATION OF INTERESTS

The authors declare no competing interests.

## DECLARATION OF AI-ASSISTED TECHNOLOGIES

During the preparation of this manuscript the authors used Microsoft Copilot to assist with language refinement and improvement of logical flow. The authors reviewed and edited all AI-generated content to ensure accuracy and integrity of the scientific work. No AI tools were used for data analysis, interpretation or general scientific content.

## SUPPLEMENTAL INFORMATION

### Supplemental Figure Legends

**Figure S1. KDM4A localizes to the basal body.**

(A-D) Representative structured illumination microscopy (SIM) immunofluorescence images of hTERT RPE-1 cells co-labeled with KDM4A (green) and Centrin-2 (red) (A-B) or γ-tubulin (red) (C-D). Nuclei were counterstained with DAPI (blue). Schematics on the right indicate the viewing angle. Scale bar, 10 μm.

(E-G) 3D single-molecule super-resolution reconstructions of KDM4A (magenta) and acetylated α-tubulin (cilia marker, green) in hTERT-RPE1 cells, shown as a merge and individual channels. Insets show corresponding diffraction-limited (DL) images. Cilium 2 is also shown in Figure 1C-1E with alternative views. Scale bars: 4 µm for cilium 2; 2 µm for cilia 3 and 4.

**Figure S2. KDM4A is required for cilia formation.**

(A) Representative immunofluorescence images of hTERT RPE-1 cells expressing siControl or siKDM4A (n=3) after 48 h of serum depletion, co-labeled with PGT (cilia marker, green) and pericentrin (basal body marker, red). Nuclei were counterstained with DAPI (blue). Scale bar, 10μm.

(B) Quantification of ciliated hTERT RPE-1 cells (y-axis) expressing siControl (blue bars) or siKDM4A (pink bars) labeled with PGT. Each point represents values from individual experiments (n=3), with >500 cells analyzed per experiment. Data are shown as mean ± S.D.; * p-values as indicated (Student’s t-test).

(C, D) Quantification of cilia length (y-axis) in hTERT RPE-1 cells expressing sgNT (blue bars) or sgKDM4A (pink bars) labeled with acetylated α-tubulin (C) or Arl13b (D). Each point represents values from individual experiments (n=3 for C; n=4 for D), with >500 cells analyzed per experiment. Data are shown as mean ± S.D.; * p-values as indicated (Welch’s t-test).

(E, F) Quantification of cilia length (y-axis) in hTERT RPE-1 cells treated with vehicle (gray bars) or JIB-04 (plum bars) labeled with acetylated α-tubulin (E) or Arl13b (F). Each point represents values from individual experiments (n=3 for each), with >500 cells analyzed per experiment. Data are shown as mean ± S.D.; * p-values as indicated (Welch’s t-test).

Figure S3. KDM4A exclusively impairs ciliogenesis among Jumonji demethylases targeted by JIB-04.

(A) Representative immunofluorescence images of hTERT RPE-1 cells expressing siControl or siKDM4B, siKDM4C, siKDM5A and siKDM6B (n=3) after 48 h of serum depletion, co-labeled with PGT (cilia marker, red) and pericentrin (basal body marker, green). Nuclei were counterstained with DAPI (blue). Scale bar, 10 μm.

(B) Quantification of ciliated hTERT RPE-1 cells (y-axis) expressing siControl (blue bar) or siKDM4B (purple bar), KDM4C (green bar), KDM5A (maroon bar) and KDM6B (gray bar), labeled with PGT. Each point represents values from individual experiments (n=3), with >500 cells analyzed per experiment. Data are shown as mean ± S.D.; * p-values as indicated (Student’s t-test).

**Figure S4. KDM4A enzymatic activity preserves cilia stability.**

(A) Representative immunofluorescence images of pre-ciliated hTERT RPE-1 cells treated with vehicle (top) or increasing doses of JIB-04 (bottom), co-labeled with acetylated α-tubulin (cilia marker, red) and pericentrin (basal body marker, green). Nuclei were counterstained with DAPI (blue). Scale bar, 10 μm.

(B) Quantification of ciliated hTERT RPE-1 cells (y-axis) treated with vehicle (gray bar) or JIB-04 (plum bars). Each point represents values from individual experiments (n=2), with >500 cells analyzed per experiment. Data are shown as mean ± S.D.; * p-values as indicated (Welch’s t-test).

**Figure S5. Live cell imaging showing destabilization of cilia in conditions of KDM4A inhibition.**

(A-B) Representative time points of two cilia visualized using live cell imaging of hTERT RPE-1 cells transfected with GFP-Arl13b to label cilia (green). Cells were pre-ciliated prior to treatment with JIB-04 (10 μM) and images collected on a Nikon W1 spinning disk microscope using a 60X oil objective.

**Figure S6. AURKA inhibition fails to rescue cilia defects associated with KDM4A inhibition.**

(A) Representative immunofluorescence images of hTERT RPE-1 cells treated with vehicle (top), Alisertib (2 μM), JIB-04 (5 μM) or a combination of JIB-04 (5 μM) and Alisertib (2 μM), co-labeled with acetylated α-tubulin (cilia marker, green) and pericentrin (basal body marker, red). Nuclei were counterstained with DAPI (blue). Scale bar, 10 μm.

(A) Representative immunofluorescence images of pre-ciliated hTERT RPE-1 cells treated with vehicle (top), Alisertib (2 μM), JIB-04 (10 μM) or a combination of JIB-04 (10 μM) and Alisertib (2 μM), co-labeled with acetylated α-tubulin (cilia marker, green) and pericentrin (basal body marker, red). Nuclei were counterstained with DAPI (blue). Scale bar, 10 μm.

**Figure S7. KDM4A regulates inter-centriolar distances.**

(A-B) Quantification of inter-centriolar distances (μm, y-axis) in pre-ciliated hTERT RPE-1 cells treated with vehicle (gray) or JIB-04 (plum) for 24 h. Each point represents values from one representative experiment (A-experiment 2; B-experiment 3), with >400 cells analyzed. * p-values as indicated (Welch’s t-test).

## METHODS

### Cell Culture

Immortalized human retinal pigmented epithelial (hTERT RPE-1) cells (gift from Dr. Gregory Pazour, University of Massachusetts Medical School, Worcester, MA) were maintained in DMEM/F-12 (ThermoFisher Scientific, Cat. #11320-082) and cultured at 37°C and 5% CO_2_. All cell lines were supplemented with 10% Fetal Bovine Serum (FBS, Sigma-Aldrich, Cat. # F2442). Cell lines used in this study were routinely monitored for mycoplasma and confirmed negative before use for experiments. In addition, human cell lines were short tandem repeat fingerprinted and validated using the Characterized Cell Line Core Facility (U.T. M. D. Anderson Cancer Center).

### Generation of Stable Cell Lines

*Kdm4a CRISPR RPE-1 cells: Kdm4a* was knocked out in hTERT RPE-1 cells using CRISPR-Cas9 gene editing. sgRNA guides (sgKdm4a) co-expressing Cas9 were purchased from GenScript. Three sgRNA constructs were co-transfected into RPE-1 cells and selected using puromycin (2 μg/ml) (ThermoFisher Scientific, Cat. # A11138-03). Single colonies were selected and validated by immunoblotting to confirm *Kdm4a* knockout. The sequences for the guides used are as follows: sgKdm4a#1- TAGATCATCAATATCGTCGT, sgKdm4a#2 – GATCTTGCGGAACTCACGAA, sgKdm4a#3 – CGGCCGGCTGAAGACCATCC.

### Transfections and Drug Treatments

*siRNAs:* siControl (Cat. # D-001810-01-20), siKDM4A (Cat.# M-004292-01-0010), siKDM4B (Cat. # L-004290-00-0010), siKDM4C (Cat. # L-004293-01-0010), siKDM5A (Cat. # L-003297-02-0010), siKDM6B (Cat. # L-023013-01-0010), were obtained as lyophilized powder from Dharmacon (Revvity) and resuspended in 250µL of 1X siRNA buffer (Dharmacon Cat. #B-002000-UB-100) to obtain a 20μM final stock. Cells at 70% confluence were transfected at 10 nm using DharmaFECT1 transfection reagent (Dharmacon, Cat. # T-2001-03) each and serum starved for 48 h to induce the formation of primary cilia.

*JIB-04 treatment:* JIB-04 (Z-isomer and E-isomer) was a kind gift of Dr. Elisabeth Martinez (U.T. Southwestern Medical School). JIB-04 was also purchased from Selleck Chemicals (Cat. # S7281) and resuspended in DMSO to obtain 100mM stocks. Cells at 95% confluence were treated with JIB-04 in serum-free media for 48 h to induce cilia formation. Alternately, cells were allowed to ciliate (48 h in serum-free media) and then treated with JIB-04 for 24 h for cilia stability assays. DMSO was used as a vehicle control. Cells were subsequently fixed with 4% paraformaldehyde (PFA) for immunofluorescence (see below) or harvested to prepare whole cell extracts for immunoblotting (see below).

*DFX-treatment:* Deferoxamine mesylate (DFX) was purchased from Sigma-Aldrich (Cat. # 252750) and was resuspended in nuclease-free water to a concentration of 100mM. Cells were plated and treated at 95% confluency with DFX at 250μM in serum-free media for 48 h (to induce cilia formation). Nuclease-free water was used as a vehicle control. Cells were subsequently fixed with 4% PFA for immunofluorescence (see below) or harvested to prepare whole cell extracts for immunoblotting (see below).

### Direct Immunofluorescence

Immunofluorescence staining was performed to visualize primary cilia in hTERT-RPE1 cells. Cells were washed with autoclaved PBS and fixed with 4% PFA (Electron Microscopy Sciences, Cat. # 15714) for 15 min at RT, followed by permeabilization with 0.5% Triton-X for 10 min, and then blocked for 1 h using 3.75% bovine serum albumin (BSA) (Sigma Aldrich, Cat. #A8412), which was used as the blocking buffer and to dilute both primary and secondary antibodies. Samples were incubated with primary antibodies overnight at 4°C. The following day, cells were washed 3 times (10 min/wash) with autoclaved PBS before application of secondary antibodies; anti-mouse Alexa Fluor 488, 546, 647 (Life Technologies, Cat. # A11001, A11003, and A21235, respectively) and anti-rabbit Alexa Fluor 488, 546, 647 (Life Technologies, Cat. # A11034, A11010, and A21245, respectively) for 1 h at RT in 3.75% BSA. Cells were washed 3 times for 10 min each with autoclaved PBS and post-fixed with 4% PFA, followed by counterstaining with DAPI (1:4000, ThermoFisher Scientific, Cat # 62248). For the co-immunofluorescence of KDM4A and Rootletin, since both antibodies were generated to the same species (rabbit), the primary and secondary antibodies were pre-incubated separately with anti-KDM4A antibody and anti-Rootletin antibody for 1.5 hours at 4°C. Following blocking and permeabilization of cells, the two individual pre-incubated antibodies (mixture of primary and secondary) were added to the cells at a 1:1 ratio for 2 h prior to washing, and counterstaining with DAPI as indicated above. Coverslips were mounted onto glass slides and visualized and imaged using a Nikon Ti2 inverted microscope with a deconvolution package (Nikon, Melville, NY, USA) at 60× magnification (secondary validation assays). Immunofluorescence images were post-acquisition deconvolved using the Nikon Elements software and further analyzed for percentage of cilia and cilia lengths using in-built modules on the software.

### Single-Molecule Super-Resolution Immunofluorescence

#### Immunolabelling

To perform dSTORM imaging of KDM4A and acetylated α-tubulin, hTERT RPE-1 cells were cultured in 8-well glass-bottom µ-slides (Ibidi, Cat. # 80827, #1.5H). After fixation with chilled 100% methanol for 10 mins, the cells were blocked with 3% (w/v) bovine serum albumin (BSA) (Sigma-Aldrich, Cat. # A2058) diluted in PBS for 1 h at RT. Subsequently, the cells were labeled with rabbit anti-KDM4A primary antibodies (Sigma-Aldrich, Cat. # HPA007610, 1:250 dilution) and mouse anti-acetylated α-tubulin (Invitrogen, 32-2700, 1:250 dilution) diluted in 1% (w/v) BSA in PBS with 0.1% (v/v) Tween 20 (Promega, Cat. # H5152) overnight at 4°C. After washing the cells three times with PBS, the cells were labeled with donkey anti-rabbit AF647 (Abcam, Cat. # ab150067, 1:1000 dilution) and goat anti-mouse CF568 (Biotium, Cat. # 20105, 1:1000 dilution) in 1% (w/v) BSA in PBS with 0.1% Tween20 for 1 h at RT. Finally, the cells were washed three times with PBS.

#### Optical Setup

All single-molecule super-resolution imaging data were obtained using an optical setup built around an inverted microscope (Olympus, IX83) equipped with a 100x, 1.5 NA objective as previously described^21^. Briefly, the samples were excited by Gaussian highly inclined and laminated optical sheet (HILO) illumination at either 560 nm or 647 nm. The fluorescence signal from the sample was spectrally separated into two paths (“green” and “red”) using a dichroic mirror before entering a two-channel 4f system. For 3D imaging with double-helix point spread functions (DH-PSFs)^20,22,23,44,45^, transmissive dielectric double-helix phase masks (Double Helix Optics, Inc., Cat. # DH1-580, DH1-670) were placed in the Fourier planes in each channel. µManager^46,47^ was used to control a piezoelectric XYZ stage and image acquisition.

#### dSTORM Imaging

For diffraction-limited imaging, the cells were imaged in an imaging buffer containing 100 mM Tris-HCl (Thermo Fisher Scientific, Cat. # J22638), 10% (w/v) glucose (Sigma-Aldrich, Cat. # 50-99-7), 2 µL/mL catalase suspension (MP Biomedicals, Cat. # 10042910), and 0.56 mg/mL glucose oxidase (Sigma Aldrich, Cat. # G2133) in ultrapure water. 100 frames were acquired at 50 ms exposure time using a low-power excitation (∼ 0.7 W/cm 2) at either 560 nm or 647 nm. For 3D dSTORM imaging, the same imaging buffer was supplemented with 429 mM β-mercaptoethanol (Sigma-Aldrich, Cat. # M6250). First, KDM4A was imaged using 647 nm Gaussian HILO excitation with an intensity of 8.2 kW/cm^2^. Then, acetylated α-tubulin was imaged using 560 nm Gaussian HILO excitation with an intensity of 2.7 kW/cm^2^. For both targets, 20,000 frames were acquired at an exposure time of 50 ms. Fluorescent beads (Invitrogen, Cat. # F8806) were added to the sample to measure the sample drift during image acquisition. For DH-PSF calibration and two-channel registration, Tetraspeck fluorescent beads (Invitrogen, Cat. # T7279) diluted 1:10 in 10% polyvinyl alcohol (Polysciences, Inc., Mowiol 4-88, 17951) in ultrapure water were immobilized onto plasma-cleaned glass coverslips. For DH-PSF calibration, z-scans of the beads were acquired with a 50 nm step size using the piezoelectric XYZ stage. For channel registration, 100 frames of the bead images were acquired at five different fields of view.

#### Image analysis

The frames acquired for diffraction-limited imaging data were summed and averaged using ImageJ. For single-molecule imaging data, localization was performed using AutoDS3D software^48^ (https://github.com/alonsaguy/One-click-image-reconstruction-in-single-molecule-localization-microscopy-via-deep-learning). In AutoDS3D, convolutional neural network (CNN) models were trained on the single-molecule images simulated using the PSF (point spread function) model generated from the DH (double-helix) phase pattern retrieved from the z-scan images of a fluorescent bead on the glass coverslip. The training datasets were simulated using the parameters that matched the experimental single-molecule data obtained in the green or red channels. For the green channel, the training data spanned a signal count in the range of 6,000 to 25,000, a baseline in the range of 80 to 150, and a readout noise standard deviation in the range of 300 to 600. For the red channel, the training parameters were a signal count in the range of 8,000 to 40,000, a baseline in the range of 100 to 300, and a readout noise standard deviation in the range of 40 to 80. The trained CNN models were then used to localize molecules in the single-molecule data of KDM4A and acetylated α-tubulin. Fiducial bead positions during the image acquisition were tracked by localizing them using the PYME plugin for DH-PSF detection and localization^49^ (https://zenodo.org/records/16879093). The localized bead trajectories were then used to correct the drift in the single-molecule data via cubic spline fitting in MATLAB, as described previously^22,23^. Two-channel registration was performed in MATLAB using a transformation matrix derived from the bead in the red channel to the bead in the green channel, as detailed in previous studies^18,21,50^. After the 2D transformation, any focal shift between channels was corrected by manually aligning the averaged z positions of the fiducial bead from the first 100 frames of each dataset. All the localization was rendered using Vutara SRX (Bruker) using point splatting with a particle size of 30 nm.

### Immunoblotting

Whole cell lysates (WCEs) were generated from cells collected in ice-cold PBS and pelleted by centrifugation at 10000 rpm at 4°C. Aspirated supernatant and pellets were flash-frozen in liquid nitrogen and stored at-80°C for future use or in cold 1× cell lysis buffer (20 mM Tris (pH 7.5), 300 mM NaCl, 1 mM EDTA, 1% NP-40). 1X complete protease inhibitor cocktail (Roche, Cat. # 04693132001), 1 mM sodium orthovanadate (Na3VO4, Sigma Aldrich, Cat. # S6508), and 1 mM PMSF (Sigma Aldrich, Cat. # P7626) were added fresh to the lysis buffer prior to use. The cellular extracts were sonicated for 10 cycles (30 seconds on and 30 seconds off per cycle) using a Diagenode Bioruptor 300 and the extracts centrifuged at maximum speed for 10 min at 4°C. The pellet was discarded, and the supernatant was collected as WCE. WCEs were subjected to the BCA-protein assay to quantify and normalize protein levels. Soluble proteins were subject to immunoblot analysis using 4-15 % SDS-PAGE gels (Bio-Rad, Cat. # 64557025) followed by transfer to PVDF membrane. Membranes were blocked for 1 h in 5% non-fat milk and incubated with primary antibodies overnight at 4°C. Membranes were subsequently washed 3 times with TBST on a shaker, and primary antibodies were tagged with horseradish peroxidase (HRP) conjugated goat anti-mouse and goat anti-rabbit secondary antibodies by incubation for 1 h at RT. Membranes were then washed three times with TBST (10 mins each) and visualization performed using ECL and ECL Prime (ThermoFisher Scientific, Cat. #32106 and Amersham, Cat. #RPN2232, respectively).

### Antibodies

Antibodies were procured from the following sources: anti-KDM4A (1:1000, Cat. # HPA007610), acetylated α-tubulin (1:4000, Clone 6-11B-1) and anti-Centrin (1:200, Clone20H5, Cat # 04-1624) were purchased from Sigma Aldrich. Anti-pericentrin (1:4000, Abcam, Cat. # ab4448), PGT (1:1000, Clone GT335, Cat. # AG-20B-0020-C100, Adipogen Life Sciences), anti-H3K36me3 (1:1000, Active Motif, Cat. # 61101), and anti-H3 (1:5000, Cell Signaling, Cat. # 4499). Anti-GAPDH (1:5000, Cat. # 10494-1-AP), anti-Arl13b (1:3000, Cat. # 17711-1-AP), and anti-Rootletin (Cat. # 55485-1-AP) antibodies were obtained from ProteinTech. HRP conjugated secondary goat anti-mouse (Cat. # 170-6516), and goat anti-rabbit (Cat. # 170-6515) antibodies were used at 1:2000 for immunoblotting and purchased from Biorad.

### Immunoprecipitation Mass Spectrometry and Co-immunoprecipitation

2 mg of WCEs were pre-cleared with 300 μg of Dyna-G beads (Invitrogen Cat. # 1003D) for 1 hour. After pre-clearing, lysates were immunoprecipitated with anti-KDM4A antibody overnight at 4°C. Fresh Dyna-G beads were added to this mixture for 1 hour the following morning. Beads were subsequently washed 5 times with lysis buffer (10 mM TRIS 7.5, 150 mM NaCl, 5 mM EDTA, 1% NP40, 1% sodium deoxycholate, 0.025% sodium dodecylsulphate with protease inhibitor, PMSF, sodium orthovanadate, and DTT) and 2 more times with PBS (samples were also saved for co-IP analysis). After removal of excess PBS, beads were sent for mass spectrometry analysis (see below). Simultaneously previously washed and saved samples were denatured and run on an 4-15% SDS-PAGE gel and transferred to a PVDF membrane for immunoblotting with anti-KDM4A and anti-Rootletin antibodies.

### Mass Spectrometry Analysis

The identification of the KDM4A protein complex followed established protocols^51^. Briefly, 1 μg of Trypsin/Lys-C was added directly to the beads for overnight digestion. The resulting peptides were purified and desalted using custom STAGE-tip columns^52^ containing 2 mg of C18 resin (3 µm, Dr. Maisch GmbH), followed by vacuum drying. Chromatographic separation was performed on an Easy-nLC 1000 system (Thermo Fisher Scientific) using a two-column setup: an in-house trap (2 cm × 100 µm i.d.) and a capillary analytical column (5 cm × 150 µm) packed with 1.9 µm Reprosil-Pur Basic C18 beads. Peptides were eluted at 800 nl/min using a 2–26% acetonitrile gradient in 0.1% formic acid. Mass spectra were acquired on an Orbitrap Fusion (Thermo Fisher Scientific) using Xcalibur v4.1. The instrument operated in data-dependent mode, selecting the 30 most intense precursor ions from a full MS scan (300–1400 m/z, 120,000 resolution) for HCD fragmentation. MS/MS spectra were captured in the ion trap at a rapid scan rate. Raw data were searched against the Human RefSeq database (Jan 2020) via the Mascot algorithm (v2.4) within the Proteome Discoverer 2.1 environment. Search criteria included a 20 ppm precursor tolerance, 0.5 Da fragment tolerance, and a maximum of two missed cleavages, considering N-terminal acetylation and methionine oxidation as variable modifications. A 1% FDR threshold was enforced using Percolator. Protein-level inference and MS1-based label-free quantification were managed by the gpGrouper algorithm^53^, which allocated peptides to gene products. Quantitative values were calculated using the iBAQ method. For statistical analysis, missing values were imputed using half of the minimum detected iBAQ intensity. Following log2 transformation, significant differences were determined via a Student’s t-test.

### RT-PCR Analysis

Total RNA was extracted from transfected and drug-treated cells using TRIzol reagent (ThermoFisher Scientific, Cat. # 15596026) and the resulting RNA cleaned using RNeasy Mini Kit (Qiagen, Cat. # 74104) according to the manufacturer’s instructions. The quality of the extracted RNA was determined using the RNA TapeStation 4200 (Agilent) and quantified using a Qubit Fluorometer (ThermoFisher Scientific). Complementary DNA (cDNA) was prepared by reverse transcription (Superscript III, Life Technologies). Gene expression was analyzed in 96-well plates by real-time quantitative PCR using specific TaqMan probes (Thermo Fisher Scientific, Waltham, MA, USA) and TaqMan Fast Universal master mix on an Applied Biosystems QuantStudio 6 Flex Real-Time PCR system (Thermo Fisher Scientific, Waltham, MA, USA). The mRNA expression was determined for AURKA, KDM4A, and GAPDH (control). For each real-time reaction, the conditions were: 95 °C for 20 min, followed by 40 cycles of 1 s at 95 °C and 20 s at 60 °C. All RT-PCR experiments were done in at least triplicate and quantified using the − ΔΔ cycle threshold (CT) method.

## Statistical Analysis

Statistical analyses were performed using Student’s t-test (two-tailed, assuming equal variance) and Welch’s t-test (two tailed, assuming unequal variance) from at least three biological replicates. The standard deviation of the mean (SD) was calculated, and p-values less than 0.05 were considered significant. All statistical analyses were performed using Prism 10 (GraphPad) and Excel (Microsoft) software.

## Notes

### Competing Interest Statement

The authors have declared no competing interest.

## REFERENCES

1. Pedersen, L.B., Schroder, J.M., Satir, P., and Christensen, S.T. (2012). The ciliary cytoskeleton. Compr Physiol 2, 779–803. 10.1002/cphy.c110043.

2. Mill, P., Christensen, S.T., and Pedersen, L.B. (2023). Primary cilia as dynamic and diverse signalling hubs in development and disease. Nat Rev Genet 24, 421–441. 10.1038/s41576-023-00587-9.

3. Anvarian, Z., Mykytyn, K., Mukhopadhyay, S., Pedersen, L.B., and Christensen, S.T. (2019). Cellular signalling by primary cilia in development, organ function and disease. Nat Rev Nephrol 15, 199–219. 10.1038/s41581-019-0116-9.

4. Ti, H., Zhang, Z., Yan, X., Hu, H., Zhang, K., Shi, S., Wu, J., Nie, H., Yuan, Z., Chen, Y., et al. (2025). Primary cilia as mechanosensors in musculoskeletal homeostasis and disease. Pharmacol Res 219, 107887. 10.1016/j.phrs.2025.107887.

5. Schneider, P., Fandrey, J., and Leu, T. (2025). Primary cilia as antennas for oxygen. Am J Physiol Cell Physiol 328, C381–C386. 10.1152/ajpcell.00298.2024.

6. Reiter, J.F., and Leroux, M.R. (2017). Genes and molecular pathways underpinning ciliopathies. Nat Rev Mol Cell Biol. 10.1038/nrm.2017.60.

7. Zhao, H., Khan, Z., and Westlake, C.J. (2023). Ciliogenesis membrane dynamics and organization. Semin Cell Dev Biol 133, 20–31. 10.1016/j.semcdb.2022.03.021.

8. Tapia Contreras, C., and Hoyer-Fender, S. (2021). The Transformation of the Centrosome into the Basal Body: Similarities and Dissimilarities between Somatic and Male Germ Cells and Their Relevance for Male Fertility. Cells 10. 10.3390/cells10092266.

9. Mansour, F., Boivin, F.J., Shaheed, I.B., Schueler, M., and Schmidt-Ott, K.M. (2021). The Role of Centrosome Distal Appendage Proteins (DAPs) in Nephronophthisis and Ciliogenesis. Int J Mol Sci 22. 10.3390/ijms222212253.

10. Pugacheva, E.N., Jablonski, S.A., Hartman, T.R., Henske, E.P., and Golemis, E.A. (2007). HEF1-dependent Aurora A activation induces disassembly of the primary cilium. Cell 129, 1351–1363. S0092-8674(07)00546-6 [pii] 10.1016/j.cell.2007.04.035.

11. Nishimura, Y., Yamakawa, D., Shiromizu, T., and Inagaki, M. (2021). Aurora A and AKT Kinase Signaling Associated with Primary Cilia. Cells 10. 10.3390/cells10123602.

12. Chowdhury, P., Powell, R.T., Stephan, C., Uray, I.P., Talley, T., Karki, M., Tripathi, D.N., Park, Y.S., Mancini, M.A., Davies, P., and Dere, R. (2018). Bexarotene - a novel modulator of AURKA and the primary cilium in VHL-deficient cells. J Cell Sci 131. 10.1242/jcs.219923.

13. Black, J.C., Van Rechem, C., and Whetstine, J.R. (2012). Histone lysine methylation dynamics: establishment, regulation, and biological impact. Mol Cell 48, 491–507. 10.1016/j.molcel.2012.11.006.

14. Gong, F., and Miller, K.M. (2019). Histone methylation and the DNA damage response. Mutat Res Rev Mutat Res 780, 37–47. 10.1016/j.mrrev.2017.09.003.

15. Young, N.L., and Dere, R. (2021). Mechanistic insights into KDM4A driven genomic instability. Biochem Soc Trans 49, 93–105. 10.1042/BST20191219.

16. Lee, D.H., Kim, G.W., Jeon, Y.H., Yoo, J., Lee, S.W., and Kwon, S.H. (2020). Advances in histone demethylase KDM4 as cancer therapeutic targets. Faseb J 34, 3461–3484. 10.1096/fj.201902584R.

17. Van Rechem, C., Black, J.C., Boukhali, M., Aryee, M.J., Graslund, S., Haas, W., Benes, C.H., and Whetstine, J.R. (2015). Lysine demethylase KDM4A associates with translation machinery and regulates protein synthesis. Cancer discovery 5, 255–263. 10.1158/2159-8290.CD-14-1326.

18. Chowdhury, P., Wang, X., Han, R.I., Motrapu, M., Boice, A.G., Nakatani, Y., Vargas-Hernandez, S., Love, J.F., Chew, C., Grimm, S.L., et al. (2025). Lysine demethylase 4A is a centrosome-associated protein required for centrosome integrity and genomic stability. Febs J. 10.1111/febs.70240.

19. Biven, E., and Wang, J.T. (2025). Mechanisms underlying centriole stability. The Journal of biological chemistry 301, 110869. 10.1016/j.jbc.2025.110869.

20. Gustavsson, A.K., Petrov, P.N., Lee, M.Y., Shechtman, Y., and Moerner, W.E. (2018). 3D single-molecule super-resolution microscopy with a tilted light sheet. Nat Commun 9, 123. 10.1038/s41467-017-02563-4.

21. Nelson, T., Vargas-Hernandez, S., Freire, M., Cheng, S., and Gustavsson, A.K. (2024). Multimodal illumination platform for 3D single-molecule super-resolution imaging throughout mammalian cells. Biomed Opt Express 15, 3050–3063. 10.1364/BOE.521362.

22. Nakatani, Y., Gaumer, S., Shechtman, Y., and Gustavsson, A.K. (2024). Long-Axial-Range Double-Helix Point Spread Functions for 3D Volumetric Super-Resolution Imaging. J Phys Chem B 128, 11379–11388. 10.1021/acs.jpcb.4c05141.

23. Saliba, N., Gagliano, G., and Gustavsson, A.K. (2023). Whole-cell multi-target single-molecule super-resolution imaging in 3D with microfluidics and a single-objective tilted light sheet. bioRxiv. 10.1101/2023.09.27.559876.

24. Cheng, S., Saliba, N., Gagliano, G., Joshi, P., and Gustavsson, A.K. (2025). Single-objective lattice light sheet microscopy with microfluidics for single-molecule super-resolution imaging of mammalian cells. bioRxiv. 10.1101/2025.07.18.665606.

25. Cheng, S., Saliba, N., Gagliano, G., Joshi, P., and Gustavsson, A.K. (2026). Single-Objective Lattice Light Sheet Microscopy with Microfluidics for Single-Molecule Super-Resolution Imaging of Mammalian Cells. ACS Photonics 13, 249–262. 10.1021/acsphotonics.5c02201.

26. Chen, Z., Zang, J., Kappler, J., Hong, X., Crawford, F., Wang, Q., Lan, F., Jiang, C., Whetstine, J., Dai, S., et al. (2007). Structural basis of the recognition of a methylated histone tail by JMJD2A. Proc Natl Acad Sci U S A 104, 10818–10823. 10.1073/pnas.0704525104.

27. Wang, L., Chang, J., Varghese, D., Dellinger, M., Kumar, S., Best, A.M., Ruiz, J., Bruick, R., Pena-Llopis, S., Xu, J., et al. (2013). A small molecule modulates Jumonji histone demethylase activity and selectively inhibits cancer growth. Nat Commun 4, 2035. 10.1038/ncomms3035.

28. Taylor-Papadimitriou, J., and Burchell, J.M. (2022). Histone Methylases and Demethylases Regulating Antagonistic Methyl Marks: Changes Occurring in Cancer. Cells 11. 10.3390/cells11071113.

29. Chen, J.V., Kao, L.R., Jana, S.C., Sivan-Loukianova, E., Mendonca, S., Cabrera, O.A., Singh, P., Cabernard, C., Eberl, D.F., Bettencourt-Dias, M., and Megraw, T.L. (2015). Rootletin organizes the ciliary rootlet to achieve neuron sensory function in Drosophila. J Cell Biol 211, 435–453. 10.1083/jcb.201502032.

30. Gray, Z.H., Honer, M.A., Ghatalia, P., Shi, Y., and Whetstine, J.R. (2025). 20 years of histone lysine demethylases: From discovery to the clinic and beyond. Cell 188, 1747–1783. 10.1016/j.cell.2025.02.023.

31. Lee, J., Kim, J.S., Cho, H.I., Jo, S.R., and Jang, Y.K. (2022). JIB-04, a Pan-Inhibitor of Histone Demethylases, Targets Histone-Lysine-Demethylase-Dependent AKT Pathway, Leading to Cell Cycle Arrest and Inhibition of Cancer Stem-Like Cell Properties in Hepatocellular Carcinoma Cells. Int J Mol Sci 23. 10.3390/ijms23147657.

32. Chen, M., Zhu, H., Li, J., Luo, D., Zhang, J., Liu, W., and Wang, J. (2024). Research progress on the relationship between AURKA and tumorigenesis: the neglected nuclear function of AURKA. Ann Med 56, 2282184. 10.1080/07853890.2023.2282184.

33. Chowdhury, P., Perera, D., Powell, R.T., Talley, T., Tripathi, D.N., Park, Y.S., Mancini, M.A., Davies, P., Stephan, C., Coarfa, C., and Dere, R. (2021). Therapeutically actionable signaling node to rescue AURKA driven loss of primary cilia in VHL-deficient cells. Sci Rep 11, 10461. 10.1038/s41598-021-89933-7.

34. Sells, T.B., Chau, R., Ecsedy, J.A., Gershman, R.E., Hoar, K., Huck, J., Janowick, D.A., Kadambi, V.J., LeRoy, P.J., Stirling, M., et al. (2015). MLN8054 and Alisertib (MLN8237): Discovery of Selective Oral Aurora A Inhibitors. ACS Med Chem Lett 6, 630–634. 10.1021/ml500409n.

35. van Hoorn, C., and Carter, A.P. (2024). A cryo-electron tomography study of ciliary rootlet organization. Elife 12. 10.7554/eLife.91642.

36. Yang, J., Adamian, M., and Li, T. (2006). Rootletin interacts with C-Nap1 and may function as a physical linker between the pair of centrioles/basal bodies in cells. Mol Biol Cell 17, 1033–1040. 10.1091/mbc.e05-10-0943.

37. Mahen, R. (2021). The structure and function of centriolar rootlets. J Cell Sci 134. 10.1242/jcs.258544.

38. Yang, J., Gao, J., Adamian, M., Wen, X.H., Pawlyk, B., Zhang, L., Sanderson, M.J., Zuo, J., Makino, C.L., and Li, T. (2005). The ciliary rootlet maintains long-term stability of sensory cilia. Molecular and cellular biology 25, 4129–4137. 10.1128/MCB.25.10.4129-4137.2005.

39. Mohan, S., Timbers, T.A., Kennedy, J., Blacque, O.E., and Leroux, M.R. (2013). Striated rootlet and nonfilamentous forms of rootletin maintain ciliary function. Curr Biol 23, 2016–2022. 10.1016/j.cub.2013.08.033.

40. Jiang, X., Ho, D.B.T., Mahe, K., Mia, J., Sepulveda, G., Antkowiak, M., Jiang, L., Yamada, S., and Jao, L.E. (2021). Condensation of pericentrin proteins in human cells illuminates phase separation in centrosome assembly. J Cell Sci 134. 10.1242/jcs.258897.

41. Bahe, S., Stierhof, Y.D., Wilkinson, C.J., Leiss, F., and Nigg, E.A. (2005). Rootletin forms centriole-associated filaments and functions in centrosome cohesion. J Cell Biol 171, 27–33. 10.1083/jcb.200504107.

42. Mahen, R. (2022). cNap1 bridges centriole contact sites to maintain centrosome cohesion. PLoS biology 20, e3001854. 10.1371/journal.pbio.3001854.

43. King, S.M., Sakato-Antoku, M., Patel-King, R.S., and Balsbaugh, J.L. (2024). The methylome of motile cilia. Mol Biol Cell 35, ar89. 10.1091/mbc.E24-03-0130.

44. Pavani, S.R., Thompson, M.A., Biteen, J.S., Lord, S.J., Liu, N., Twieg, R.J., Piestun, R., and Moerner, W.E. (2009). Three-dimensional, single-molecule fluorescence imaging beyond the diffraction limit by using a double-helix point spread function. Proc Natl Acad Sci U S A 106, 2995–2999. 10.1073/pnas.0900245106.

45. Gustavsson, A.K., Petrov, P.N., and Moerner, W.E. (2018). Light sheet approaches for improved precision in 3D localization-based super-resolution imaging in mammalian cells [Invited]. Opt Express 26, 13122–13147. 10.1364/OE.26.013122.

46. Edelstein, A., Amodaj, N., Hoover, K., Vale, R., and Stuurman, N. (2010). Computer control of microscopes using microManager. Curr Protoc Mol Biol Chapter 14, Unit14 20. 10.1002/0471142727.mb1420s92.

47. Edelstein, A.D., Tsuchida, M.A., Amodaj, N., Pinkard, H., Vale, R.D., and Stuurman, N. (2014). Advanced methods of microscope control using muManager software. J Biol Methods 1. 10.14440/jbm.2014.36.

48. Saguy, A., Xiao, D., Narayanasamy, K.K., Nakatani, Y., Saliba, N., Gagliano, G., Gustavsson, A.K., Heilemann, M., and Shechtman, Y. (2025). One-click reconstruction in single-molecule localization microscopy via experimental parameter-aware deep learning. Npj Imaging 3, 61. 10.1038/s44303-025-00123-w.

49. Barentine, A.E.S., Balaji, A., and Moerner, W.E. (2025). Efficient Double Helix Detection with Steerable Filters. bioRxiv. 10.1101/2025.08.14.670427.

50. Kanie, T., Liu, B., Love, J.F., Fisher, S.D., Gustavsson, A.K., and Jackson, P.K. (2025). A hierarchical pathway for assembly of the distal appendages that organize primary cilia. Elife 14. 10.7554/eLife.85999.

51. Lee, C.S., Jung, S.Y., Yee, R.S.Z., Agha, N.H., Hong, J., Chang, T., Babcock, L.W., Fleischman, J.D., Clayton, B., Hanna, A.D., et al. (2023). Speg interactions that regulate the stability of excitation-contraction coupling protein complexes in triads and dyads. Commun Biol 6, 942. 10.1038/s42003-023-05330-y.

52. Rappsilber, J., Ishihama, Y., and Mann, M. (2003). Stop and go extraction tips for matrix-assisted laser desorption/ionization, nanoelectrospray, and LC/MS sample pretreatment in proteomics. Anal Chem 75, 663–670. 10.1021/ac026117i.

53. Saltzman, A.B., Leng, M., Bhatt, B., Singh, P., Chan, D.W., Dobrolecki, L., Chandrasekaran, H., Choi, J.M., Jain, A., Jung, S.Y., et al. (2018). gpGrouper: A Peptide Grouping Algorithm for Gene-Centric Inference and Quantitation of Bottom-Up Proteomics Data. Mol Cell Proteomics 17, 2270–2283. 10.1074/mcp.TIR118.000850.

