## Supplementary figures and images for "Lysine Demethylase 4A (KDM4A) Maintains Basal Body Architecture and Protects Against Ciliary Destabilization"

### Manuscript

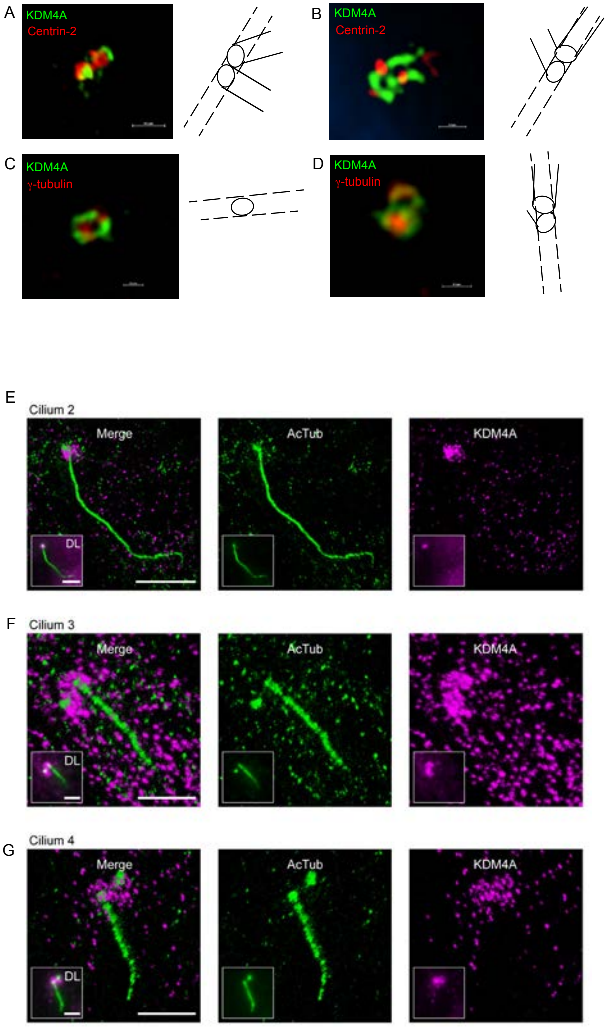

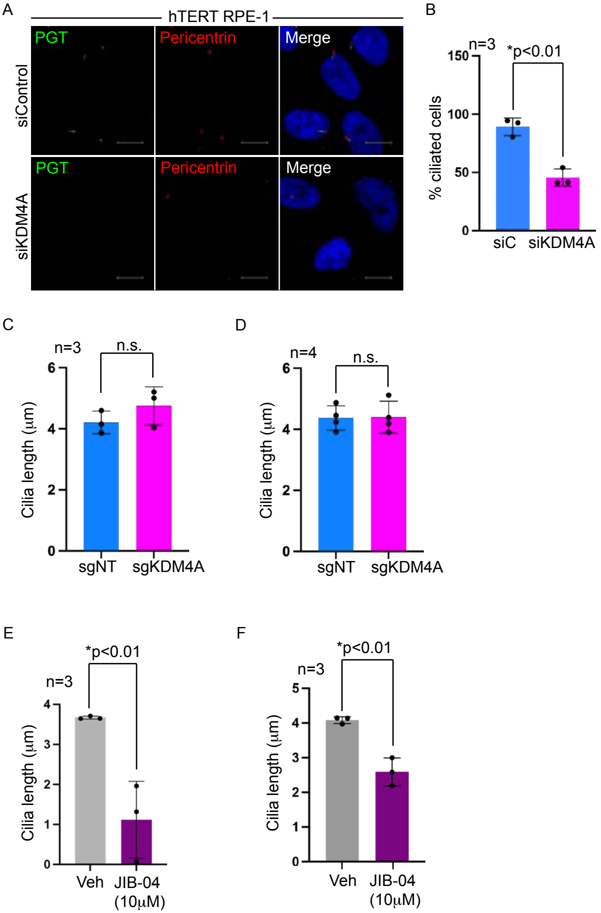

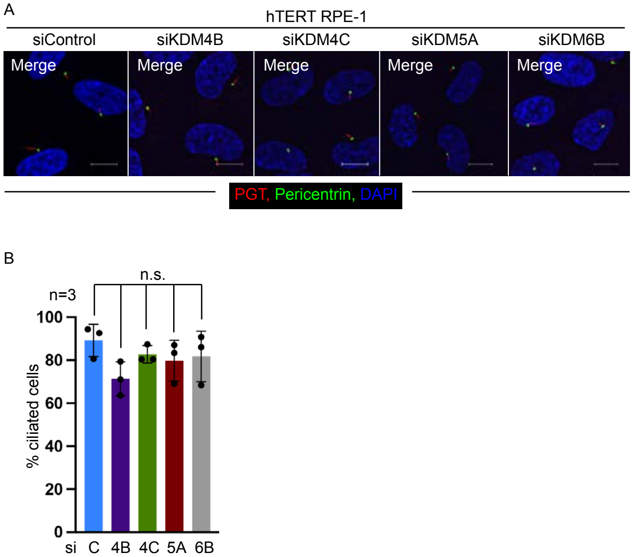

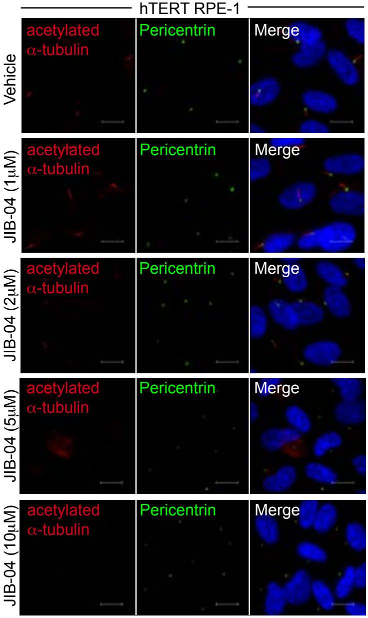

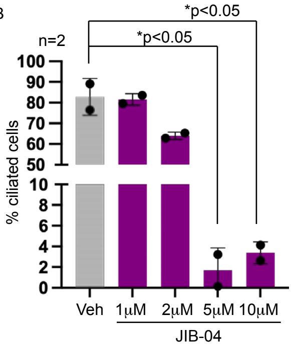

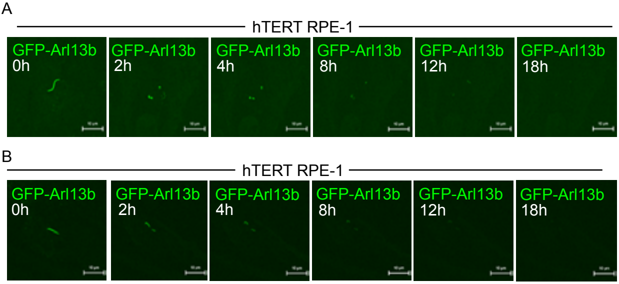

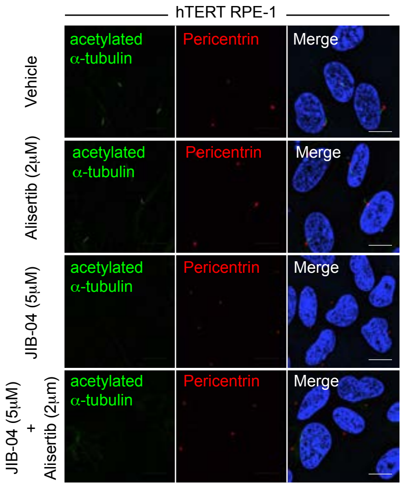

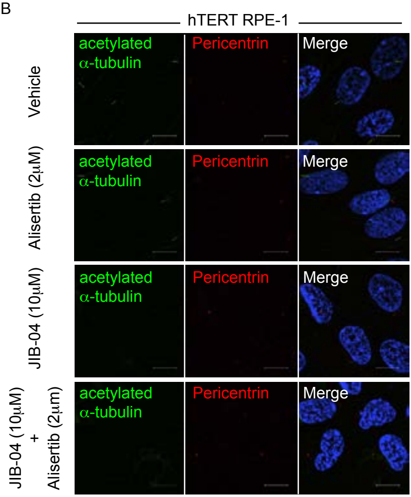

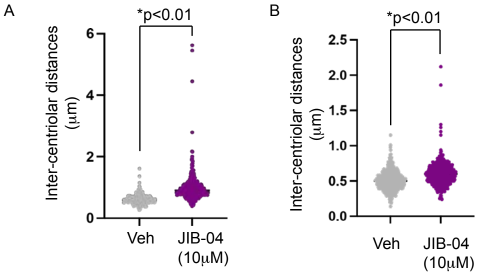
